# Anterior cingulate cortex represents action-state predictions and causally mediates model-based reinforcement learning in a two-step decision task

**DOI:** 10.1101/126292

**Authors:** Thomas Akam, Ines Rodrigues-Vaz, Ivo Marcelo, Xiangyu Zhang, Michael Pereira, Rodrigo Freire Oliveira, Peter Dayan, Rui M. Costa

**Affiliations:** Champalimaud Neuroscience Program, Champalimaud Centre for the Unknown, Lisbon, Portugal; Department of Experimental Psychology, Oxford University, Oxford, UK; Department of Neuroscience and Neurology, Zuckerman Mind Brain Behavior Institute, Columbia University, New York, NY, USA; Department of Psychiatry, Erasmus MC University Medical Center, Rotterdam, 3015 GD, The Netherlands; RIKEN-MIT Center for Neural Circuit Genetics at the Picower Institute for Learning and Memory, Department of Biology and Department of Brain and Cognitive Sciences. Massachusetts Institute of Technology, Cambridge, Massachusetts, USA; Gatsby Computational Neuroscience Unit, UCL, London, UK; Max Planck Institute for Biological Cybernetics, Tübingen, Germany; University of Tübingen, Germany

## Abstract

The anterior cingulate cortex (ACC) is implicated in learning the value of actions, but it remains poorly understood whether and how it contributes to model-based mechanisms that use action-state predictions and afford behavioural flexibility. To isolate these mechanisms, we developed a multi-step decision task for mice in which both action-state transition probabilities and reward probabilities changed over time. Calcium imaging revealed ramps of choice-selective neuronal activity, followed by an evolving representation of the state reached and trial outcome, with different neuronal populations representing reward in different states. ACC neurons represented the current action-state transition structure, whether state transitions were expected or surprising, and the predicted state given chosen action. Optogenetic inhibition of ACC blocked the influence of action-state transitions on subsequent choice, without affecting the influence of rewards. These data support a role for ACC in model-based reinforcement learning, specifically in using action-state transitions to guide subsequent choice.

**Highlights:**

- A novel two-step task disambiguates model-based and model-free RL in mice.
- ACC represents all trial events, reward representation is contextualised by state.
- ACC represents action-state transition structure, predicted states, and surprise.
- Inhibiting ACC impedes action-state transitions from influencing subsequent choice.

## Introduction

The anterior cingulate cortex (ACC) is a critical contributor to reward guided decision making (Rushworth and Behrens, 2008; Heilbronner and Hayden, 2016). ACC neurons encode diverse decision variables (Cai and Padoa-Schioppa, 2012; Ito et al., 2003; Matsumoto et al., 2003; Sul et al., 2010), and the structure has been particularly associated with action reinforcement (Hadland et al., 2003; Kennerley et al., 2006; Rudebeck et al., 2008). However, instrumental learning about the value of actions is not a unitary phenomenon, but rather is thought to be mediated by partly parallel control systems, model-based and model-free, that use different computational principles to evaluate choices (Balleine and Dickinson, 1998; Daw et al., 2005; Dolan and Dayan, 2013). Despite suggestive evidence of ACC’s involvement in model-based reinforcement (Daw et al., 2011; Cai and Padoa-Schioppa, 2012; Karlsson et al., 2012; O’Reilly et al., 2013; Doll et al., 2015; Huang et al., 2020), studies designed to specifically test this are lacking.

To investigate the ACC’s role, we need a clear articulation of these parallel systems and a paradigm that allows their contributions to be distinguished. The former stems from the venerable dissociation between habitual and goal-directed control (Balleine and Dickinson, 1998; Daw et al., 2005). Well-practiced actions in familiar environments are controlled by a habitual system, thought to employ model-free reinforcement learning (RL) (Sutton and Barto, 1998). This uses reward prediction errors to cache preferences between actions. However, when the environment or motivational state changes, model-free preferences can become out of date, and actions are instead controlled by a goal-directed system believed to utilise model-based RL (Sutton and Barto, 1998). This learns a predictive model of the consequences of actions, i.e. the states and rewards they immediately lead to, and evaluates options by simulating or otherwise estimating their resulting long-run values. This dual controller approach is beneficial because model-free and model-based RL have complementary strengths, the former allowing quick and computationally cheap decision making at the cost of slower adaptation to changes in the environment, the latter flexible and efficient use of new information at the cost of computational effort and decision speed.

For a paradigm that might distinguish between these systems, we started with the recent class of multi-step decision tasks (Daw et al., 2011; Simon and Daw, 2011; Huys et al., 2012). Canonically, on each trial, subjects traverse states in a decision tree to reach rewards, often with ongoing changes in the state transition and/or reward probabilities to force continuous learning and surface differences between flexible and inflexible decision-making processes. The so-called two-step task (Daw et al., 2011) is perhaps the most popular, with variants used to probe mechanisms of model-based RL (Daw et al., 2011; Wunderlich et al., 2012; Smittenaar et al., 2013; Doll et al., 2015) and arbitration between controllers (Keramati et al., 2011; Lee et al., 2014; Doll et al., 2016), and to identify behavioural differences in psychiatric disorders (Sebold et al., 2014; Voon et al., 2015; Gillan et al., 2016). Versions of the two-step task for rats (Miller et al., 2017; Dezfouli and Balleine, 2017; Hasz and Redish, 2018; Groman et al., 2019) and monkeys (Miranda et al., 2019) have recently been developed.

However, we have shown that with the sort of extensive experience on two-step tasks necessary for investigations with animals, subjects can, in principle, acquire a sophisticated, memory-based, representation of a latent state of the environment which confounds model-free and model-based planning (Akam et al., 2015). This would limit our ability to determine the ACC’s specific contributions.

Here, we report a novel murine two-step task designed to avoid this confound, and apply the task to probe the involvement of ACC in model-based and model-free control. The new task induces unsignalled structural changes in the decision-tree that complicate the use of latent state based strategies, whilst still permitting conventional model-based planning. We show that mice readily learn this task and show behaviour consistent with a mixture of model-based and model-free RL.

Calcium imaging of ACC neurons whilst animals performed the task revealed that different populations participated across the different stages of each trial, representing all trial events, but with a stronger representation of states reached in the decision tree than rewards obtained, and different neurons representing reward in different states. Additionally, the ACC represented a set of variables required for model-based RL, including the current configuration of the action-state transition probabilities (i.e., the probabilities of transitions in the decision tree), the actual predicted state given chosen action, and whether observed state transitions were expected or surprising given current knowledge of the tree. Consistent with this, single-trial optogenetic inhibition of ACC selectively disrupted the influence of action-state transitions on subsequent choice, while sparing the influence of rewards. Accordingly, the strength of the effect of ACC inhibition for each individual subject was closely correlated with the degree to which that subject used model-based RL to solve the task.

## Results

### A novel two-step task with transition probability reversals

As in the original two-step task (Daw et al., 2011), our task consisted of a choice between two ‘first-step’ actions which led probabilistically to one of two ‘second-step’ states where reward could be obtained. Unlike the original task, in each second-step state there was a single action rather than a choice between two actions. In the original task, the stochasticity of state transitions and reward probabilities causes both model-based and model-free control to obtain rewards at a rate negligibly different from random choice at the first-step (Akam et al., 2015; Kool et al., 2016). To promote task engagement, we increased the contrast between good and bad options by using a block-based reward probability distribution rather than the random walks used in the original, and increased the probability of common relative to rare state transitions. The final and most significant structural change was the introduction of reversals in the transition probabilities mapping the first-step actions to the second-step states. This was done to prevent habit like strategies consisting of mappings from the second-step state where rewards have recently been obtained to specific actions at the first step (Akam et al., 2015). In supplementary results we directly compare versions of the task with fixed and changing action-state transition probabilities (Figure S1); subject’s behaviour was radically different in each, suggesting that they recruit different behavioural strategies.

We implemented the task using a set of four nose-poke ports: top and bottom ports in the centre, flanked by left and right ports (Figure 1A). Each trial started with the central ports lighting up, requiring a choice between top and bottom ports. The choice of a central port led probabilistically to a ‘left-active’ or ‘right-active’ state, in which respectively the left or right port was illuminated. The subject then poked the illuminated left or right side port to gain a probabilistic water reward (Figure 1A,B). A 1 second inter-trial interval started when the subject exited the side port.

**Figure 1.**
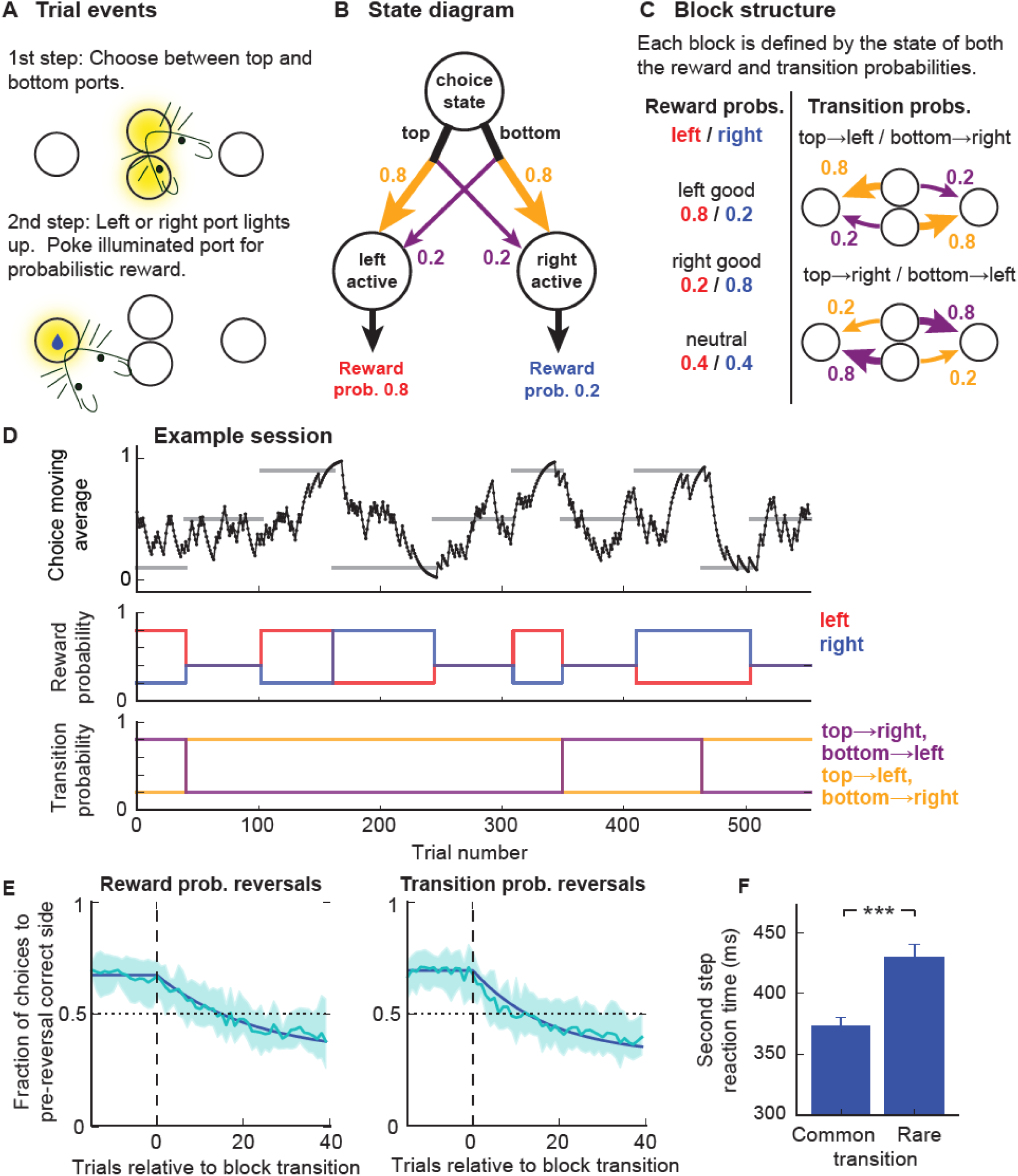
Two-step task with transition probability reversals. **A)** Diagram of apparatus and trial events. **B)** State diagram of task. Reward and transition probabilities are indicated for one of the six possible block types. **C)** Block structure, left side shows the three possible states of the reward probabilities, right side shows the two possible states of the transition probabilities. **D)** Example session: Top panel - Exponential moving average (tau = 8 trials) of choices. Horizontal grey bars show blocks, with correct choice (top, bottom or neutral) indicated by y position of bars. Middle panel – reward probabilities in left-active (red) and right-active (blue) states. Bottom panel – Transition probabilities linking first-step actions (top, bottom pokes) to second step states (left/right active). **E)** Choice probability trajectories around reversals. Pale blue line – average trajectory, dark blue line – exponential fit, shaded area – cross-subject standard deviation. Left panel - reversals in reward probability, right panel – reversals in transition probabilities. **F)** Second step reaction times following common and rare transitions - i.e. the time between the first step choice and side poke entry. *** indicates P < 0.001 Error bars show cross-subject SEM.

Both the transition probabilities linking the first-step actions to the second-step states, and the reward probabilities in each second-step state, changed in blocks. There were three possible states of the reward probabilities for the *left*/*right* ports: respectively *good/bad, neutral/neutral* and *bad/good* (Figure 1C), where good/neutral/bad reward probabilities were 0.8/0.4/0.2. There were two possible states of the transition probabilities: *top→ left / bottom→ right* and *top→ right / bottom→ left* (Figure 1C), where e.g. *top→ right* indicates the top port commonly (0.8 of trials) lead to the right port and rarely (0.2 of trials) to the left port (Figure 1C). At block transitions, the reward and/or transition probabilities changed (see figure S2 for block transition structure). Reversals in which first-step action (top or bottom) had higher reward probability could therefore occur due to reversals in either the reward or transition probabilities. Block transitions were triggered when an exponential moving average (tau = 8 trials) of the proportion of correct choices reached a threshold of 0.75, with a delay of 20 trials between threshold crossing and the reversal occurring to allow an unbiased assessment of performance at the end of blocks. This resulted in block lengths of 63.6 ± 31.7 (mean ± SD) trials.

Subjects learned the task in 3 weeks with minimal shaping and performed an average of 576 ± 174 (mean ± SD) trials per day thereafter. Our behavioural dataset used data from day 22 of training onward (n=17 mice, 400 sessions). Subjects tracked which first-step action had higher reward probability (Figure 1D,E), choosing the correct option at the end of non-neutral blocks with probability 0.68 ± 0.03 (mean ± SD). Choice probabilities adapted faster following reversals in the action-state transition probabilities (exponential fit tau = 17.6 trials), compared with reversals in the reward probabilities (tau = 22.7 trials, P = 0.009, bootstrap test, Figure 1E). Reaction times to enter the second step port were faster following common than rare transitions (P = 2.8 x 10^−8^, paired t-test) (Figure 1F).

### Disambiguating model-based and model-free strategies in the two-step task with transition probability reversals

To dissociate the contribution of model-based and model-free RL to subjects’ behaviour we looked at the granular structure of how events on each trial affected subsequent choices. The simplest such analysis examines the so-called stay probabilities of repeating the first-step choice for the four possible combinations of transition (common or rare) and outcome (rewarded or not) (Figure 2A,B). We quantified how the state transition, trial outcome, and their interaction predicted stay probability using a logistic regression analysis, with additional predictors to capture choice bias and correct for cross trial correlations which can otherwise can give a misleading picture of how trial events influence subsequent choice (Akam et al., 2015). Positive loading on the outcome predictor indicated that receiving reward was reinforcing (i.e. predicted staying) (P < 0.001, bootstrap test). Positive loading on the transition predictor indicated that experiencing common transitions was also reinforcing (P < 0.001). Loading on the transition-outcome interaction predictor was not significantly different from zero (P = 0.79).

**Figure 2.**
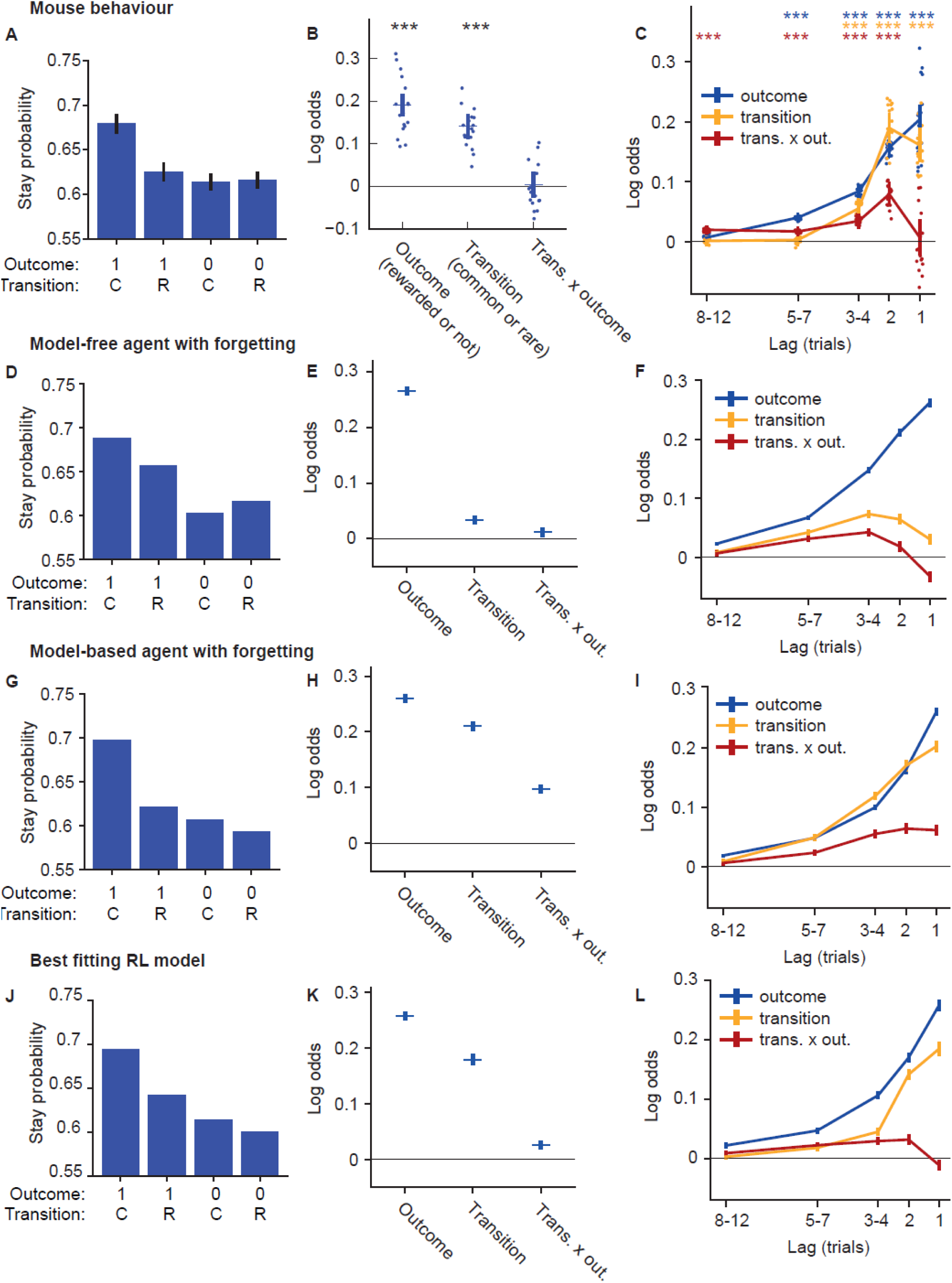
Stay probability and logistic regression analyses. **A-C)** Mouse behaviour. **A)** Stay probability analysis showing the fraction of trials the subject repeated the same choice following each combination of trial outcome (rewarded (1) or not (0)) and transition (common (C) or rare (R)). Error bars show cross-subject SEM. **B)** Logistic regression model fit predicting choice as a function of the previous trial’s events. Predictor loadings plotted are; *outcome* (repeat choices following rewards), *transition* (repeat choices following common transitions) and *transition-outcome interaction* (repeat choices following rewarded common transition trials and non-rewarded rare transition trials). Error bars indicate 95% confidence intervals on the population mean, dots indicate maximum a posteriori (MAP) subject fits. **C)** Lagged logistic regression model predicting choice as a function of events over the previous 12 trials. Predictors are as in **B. D-F)** As **A-C** but for data simulated from a model-free RL agent with forgetting and multi-trial perseveration. **G-I)** As **A-C** but for data simulated from a model-based RL agent with forgetting and multi-trial perseveration. Parameters for RL model simulations were obtained by fits of the RL models to the mouse behavioural data

The absence of a transition-outcome interaction has been used in the original two-step task (Daw 2011) to suggest that behaviour is model-free. However, we have shown (Akam et al. 2015) that this depends on the subjects not learning the transition probabilities from the experienced transitions. Such fixedness is reasonable for the original task, where transition probabilities are fixed and known to be so by the human subjects, but not for the task described here. Our analysis (Akam et al. 2015) suggests that when model-learning is included, loading in the logistic regression analysis for a model-based strategy decreases for the interaction predictor and increases for the outcome and transition predictors.

To understand the implications of this for our task better, we simulated the behaviour of a model-based and a model-free RL agent, with the parameters of both fit to the behavioural data, and ran the logistic regression analysis on data simulated from both models (Figure 2D-I). The RL agents used in these simulations included forgetting about actions not taken and states not visited, as RL model comparison indicated this greatly improved fits to mouse behaviour (see below & supplementary results). Data simulated from a model-free agent showed a large loading on the outcome predictor (i.e. rewards were reinforcing), but little loading on the transition predictor or transition-outcome interaction predictors (Figure 2E). By contrast, data simulated from the model-based agent showed a large loading on both outcome and transition predictors (i.e. both rewards and common transitions were reinforcing) (Figure 2H), and a smaller loading on the interaction predictor. Therefore, in our data the transition predictor loaded closer to the model-based strategy and the interaction predictor loaded closer to the model-free strategy.

The above analysis only considers the influence of the most recent trial’s events on choice. However, the slow time course of adaptation to reversals (Figure 1E) indicates that choices must be influenced by a longer trial history. To better understand these long-lasting effects, we used a lagged regression analysis assessing how the current choice was influenced by past transitions, outcomes and their interaction (Figure 2C). Predictors were coded such that a positive loading on e.g. the outcome predictor at lag *x* indicates that reward on trial *t* increased the probability of repeating the trial *t* choice at trial *t* + *x*. Past outcomes significantly influenced current choice up to lags of 7 trials, with a smoothly decreasing influence at larger lags. Past state transitions influenced the current choice up to lags of 4 trials with, unexpectedly, a somewhat larger influence at lag 2 compared to lag 1. Also unexpectedly, although the transition-outcome interaction on the previous trial did not significantly influence the current choice, the interaction at lag 2 and earlier did, with the strongest effect at lag 2.

To understand how these patterns relate to RL strategy, we analysed the behaviour of model-based and model-free agents using the lagged regression (Figure 2F,I). Both strategies showed a smoothly decreasing influence of trial outcome with increasing lag, similar to that observed in the data. Both strategies showed a positive loading on the transition predictor across the trial history, but this was much stronger at recent trials for the model-based strategy, similar to that observed in the data, though with a more gradual decay with increasing lag. Both strategies showed a positive loading on the transition outcome interaction predictor for earlier trials but diverged at recent trials, with the model-based strategy showing a small positive loading and the model-free a small negative loading. These data suggest that the strong influence of recent common/rare state transitions in the mouse behaviour is not consistent with a model-free strategy, however the mouse behaviour does not look like a simple mixture of model-based and model-free, suggesting the presence of additional features.

To understand how behaviour diverged from these models, we performed an in-depth model comparison, detailed in supplementary results. Here, we summarise the principal findings. As with human behaviour on the original task, the best fitting model used a mixture of model-based and model-free control. However, model comparison indicated additional features not typically used in models of two-step task behaviour: forgetting about values and state transitions for not-chosen actions, perseveration effects spanning multiple trials, and representation of actions both at the level of the choice they represent (e.g. top port) and the motor action they require (e.g. left port→top port). These are discussed in detail in the supplementary results. Taken together, the additional features substantially improved fit quality (Δ iBIC = 11018) over the model which lacked them (Figure S3). Data simulated from the best fitting RL model better matched mouse behaviour (Figure 2 J-L), with positive loading on the outcome and transition predictors and minimal loading on the interaction predictor (Figure 2J) at trial -1, but positive loading on the interaction predictor at trial -2 and earlier (Figure 2L).

These data indicate that the novel task recruits both model-based and model-free reinforcement learning mechanisms, providing a tool for mechanistic investigation into more cognitive aspects of decision making in the mouse.

### ACC activity represents all trial events, emphasises choices and states, contextualises rewards

To understand how ACC represented two-step task behaviour, we expressed GCaMP6f in ACC neurons under the CaMKII promotor (to target pyramidal neurons) and imaged calcium activity through a gradient refractive index (GRIN) lens using a miniature fluorescence microscope (n=4 mice, 21 sessions, 2385 neurons) (Ghosh et al., 2011). Constrained non-negative matrix factorisation for endoscope data (CNMF-E) (Zhou et al., 2018) was used to extract activity traces for individual neuron from the microscope video (Figure 3B). All subsequent analyses used the deconvolved activity inferred by CNMF-E. Activity was sparse, with an average event rate of 0.12Hz across the recorded population (Figure 3C). We aligned activity to the same events across trials by time-warping (see Methods) the interval between the first-step choice and second-step port entry (labelled ‘outcome’ in figures as this is when outcome information becomes available) to match the median interval. Activity prior to choice and following outcome was not time-warped. Different populations of neurons participated at different time-points across the trial (Figure 3D). Many ACC neurons ramped up activity over the 1000ms preceding the first step-choice, peaking at choice time and being largely silent following trial outcome. Other neurons were active in the period between choice and outcome, and yet others were active immediately following trial outcome.

**Figure 3.**
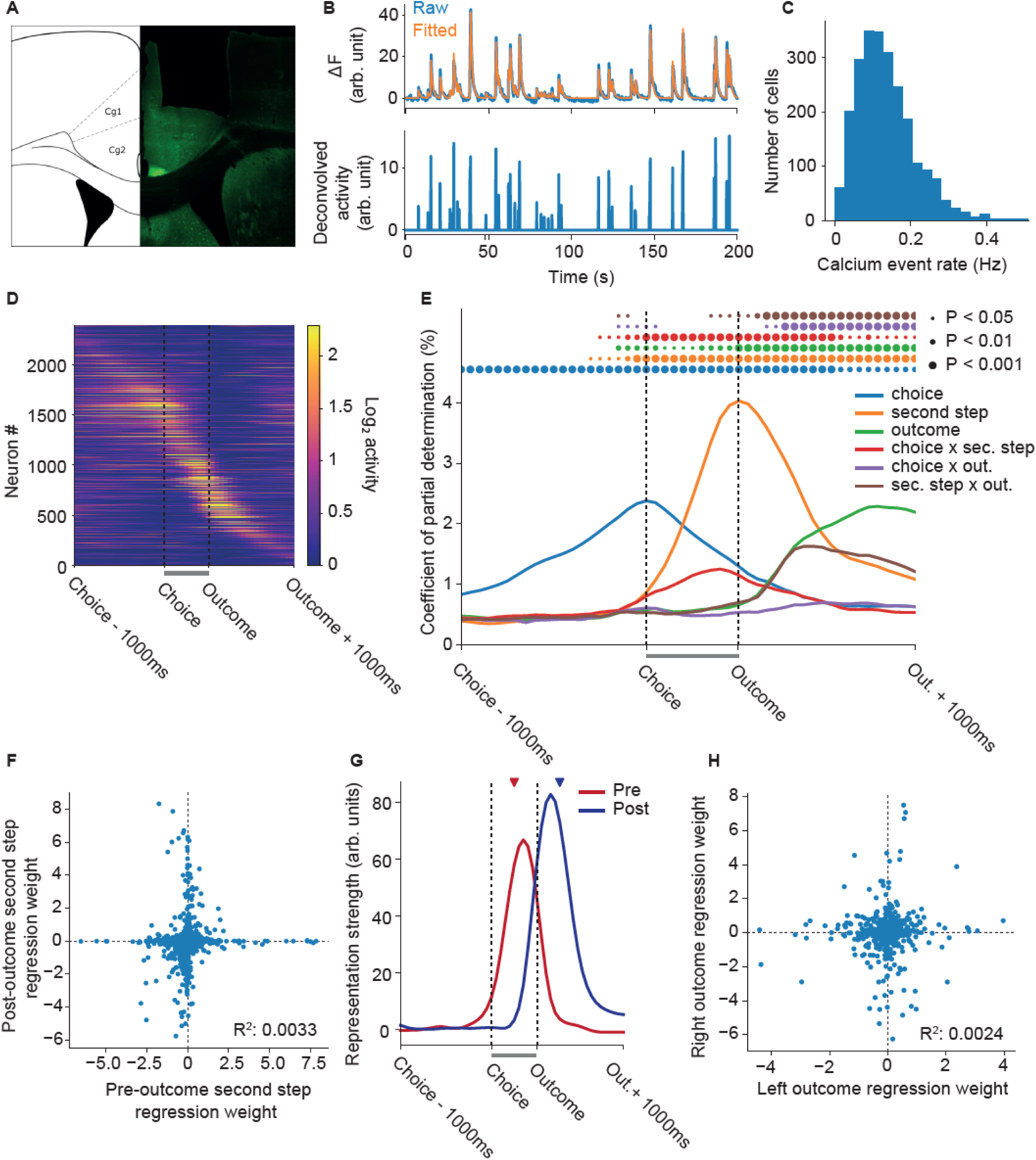
Two-step ACC calcium imaging. **A)** Example GRIN lens placement in ACC. **B)** Fluorescence signal from a neuronal ROI identified by CNMF-E (top panel – blue) and fitted trace (orange) due to the inferred deconvolved neuronal activity (bottom panel). **C)** Histogram showing the distribution of average event rates across the population of recorded neurons. Events were defined as any video frame on which the inferred activity was non-zero. **D)** Average trial aligned activity for all recorded neurons, sorted by the time of peak activity. No normalisation was applied to the activity. The grey bars under **D, E, G** between choice and outcome indicate the time period that was warped to align trials of different duration. **E)** Regression analysis predicting activity on each trial from a set of predictors coding the *choice* (top or bottom), *second step* (left or right), *outcome* (rewarded or not) that occurred on each trial, and their interactions. Lines show the population coefficient of partial determination (CPD) as a function of time relative to trial events. Circles indicate where CPD is significantly higher than expected by chance, assessed by permutation test with Benjamini–Hochberg correction for comparison at multiple time points. **F)** Representation of the second-step state before and after the trial outcome. Points show *second step* predictor loadings for individual neurons at a time-point halfway between choice and outcome (x-axis) and a time-point 250ms after trial outcome (y-axis). **G)** Time-course of pre- and post-outcome representations of second step state, obtained by projecting the second step predictor loadings at each time-point onto the pre- and post-outcome second step representations. The red and blue triangles indicate the timepoints used to define the projection vectors. **H)** Representation of trial outcomes (reward or not) obtained at the left and right poke. Points show predictor loadings for individual neurons 250ms after trial outcome in a regression analysis where outcomes at the left and right poke were coded by separate predictors.

To identify how activity represented events on the current trial, we used a linear regression predicting the activity of each neuron at each time-point as a function of the choice (top or bottom), second-step state (left or right) and outcome (rewarded or not) that occurred on the trial, as well as the interactions between these events. This and later analyses only included sessions where we had sufficient coverage of all trial types (n=3 mice, 11 sessions, 1314 neurons), as in some imaging sessions with few blocks and trials there was no coverage of trial types that occur infrequently in those blocks. We evaluated the population coefficient of partial determination, i.e. the fraction of variance across the population uniquely explained by each predictor, as a function of time relative to trial events (Figure 3E). Representation of choice ramped up smoothly over the second preceding the choice, then decayed smoothly until approximately 500ms after trial outcome. Representation of second-step state increased rapidly following the choice, peaked at second-step port entry, then decayed over the second following the outcome, and was the strongest represented trial event.

As largely distinct populations of neurons were active before and after trial outcome (Figure 3D), we asked whether the representation of second-step state was different at these two time-points by plotting the second-step state regression weights for each neuron at a time-point mid-way between choice and outcome (which we term the pre-outcome representation of second step state) against the weighs 250ms after outcome (the post-outcome representation) (Figure 3F). These pre- and post-outcome representations were uncorrelated (R^2^ = 0.0033), and neurons that were strongly tuned at one time point typically had little selectivity at the other, indicating that although second-step state was strongly represented at both times, the representations were orthogonal and involved different populations of neurons. To assess how these two representations evolved over time, we projected the regression weights for second-step state at each time-point onto the pre- and post-outcome second-step representations - i.e. onto the regression weights for second step state at these two timepoints (Figure 3G), using cross validation to give an unbiased time-course estimates. The pre-outcome representation of second step state peaked shortly before second-step port entry and decayed rapidly afterwards, while the post outcome representation peaked shortly after trial outcome and persisted for ∼500ms.

Representation of the trial outcome ramped up following receipt of outcome information (Figure 3E), accompanied by an initially equally strong representation of the interaction between trial outcome and second-step state. This interaction indicates that the representation of trial outcome depended strongly on the state in which the outcome was received. To assess this in more detail we ran a version of the regression analysis with separate predictors for outcomes received at the left and right ports, and plotted the left and right outcome regression weights 250ms after outcome against each other (Figure 3H). Representations of trial outcome obtained at the left and right port were orthogonal (R^2^ = 0.0024), indicating that although ACC carried information about reward, reward representations were specific to the state where the reward was received.

The evolving representation of trial events can be visualised by projecting the average neuronal activity for each trial type (defined by choice, second-step state and outcome) into the low dimensional space which captures the greatest variance between different trial types (see methods) (Figure 4). The first 3 principal components (PCs) of this space were dominated by representation of choice and second-step state (Figure 4A,B), with different trial outcomes being most strongly differentiated in PCs 4 and 5 (Figure 4C). Prior to the choice, trajectories diverged along an axis capturing choice selectivity (PC2). Following the choice, trajectories for different second-step states diverged first along one axis (PC3) then along a second axis (PC1), confirming that two orthogonal representations of second-step state occur in a sequence spanning the time period from choice through trial outcome.

**Figure 4.**
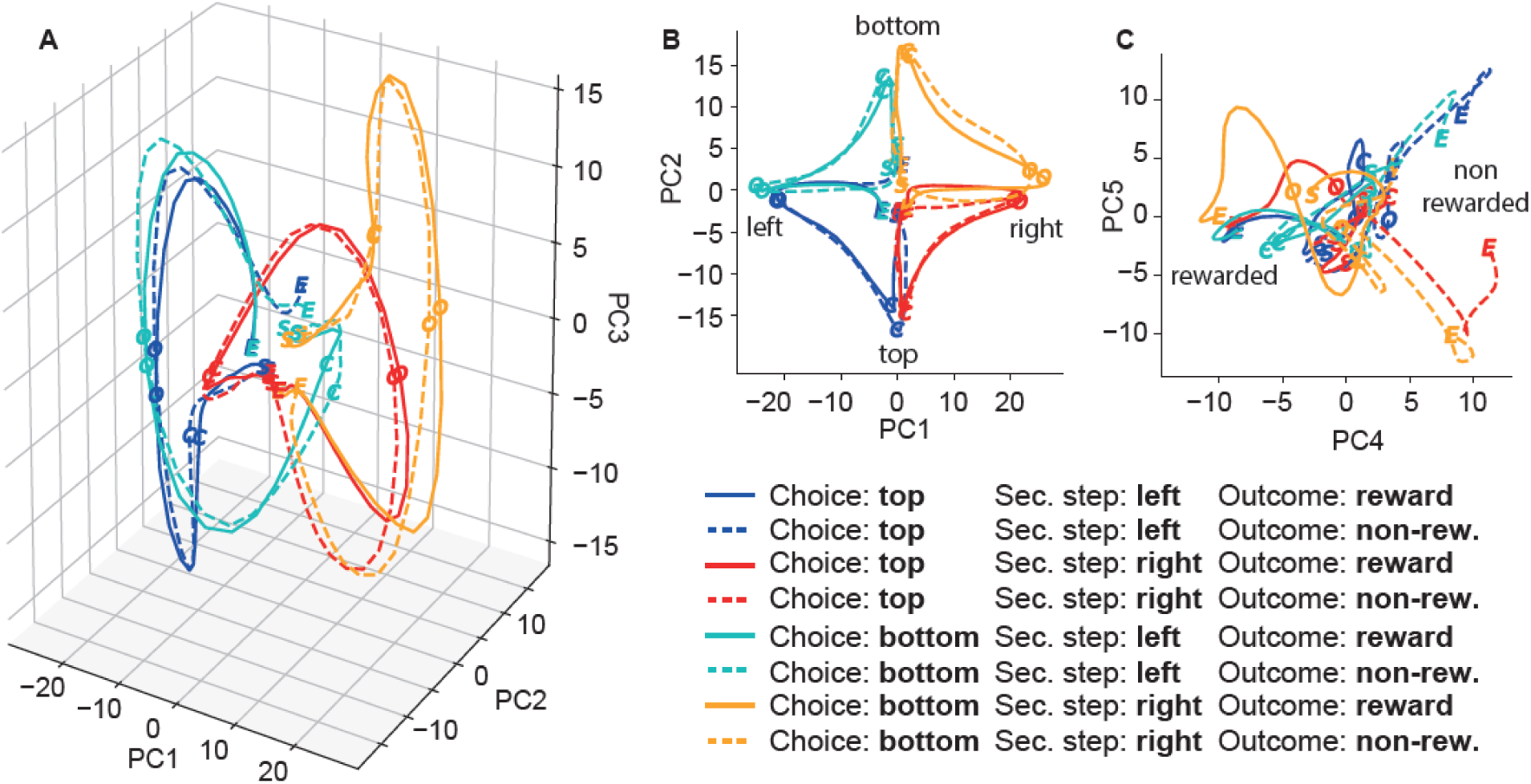
Population activity trajectories. Projection of the average population activity for different trial types into the low dimensional space which captures the most variance between trial types. Trial types were defined by the 8 combinations of choice, second-step and trial outcome. Letters on the trajectories indicate the trajectory start (*S* - 1000ms before choice), the choice (*C*), outcome (*O*) and trajectory end (*E* – 1000ms after outcome). **A**) 3D plot showing projections onto first 3 principal components. **B)** Projection onto PCs 1 and 2 which represent second-step and choice respectively. **C)** Projection onto PCs 4 and 5 which differentiate trial outcomes.

### ACC represents model-based decision variables

Model-based reinforcement learning uses predictions of the specific consequences of action, i.e. the states that actions lead to, to compute their values. Therefore if ACC implements model-based computations on this task, we expect to see representation of the current state of the transition probabilities linking first-step actions second-step states, predictions of the second-step state that will be reached given the chosen action, and surprise signals if the state that is actually reached does not match these expectations.

We therefore asked how ACC activity was affected by the changing transition probabilities mapping the first-step actions to second-step states, and reward probabilities in the second-step states. Due to the limited number of blocks that subjects performed in imaging sessions, we performed separate regression analyses for sessions where we have sufficient coverage of the different states of the transition probabilities (Figure 5A, n=3 mice, 5 sessions, 589 neurons) and reward probabilities (Figure 5B, n=3 mice, 10 sessions, 1152 neurons). These analyses predicted neuronal activity as a function of events on the current trial, the state of the transition or reward probabilities respectively, and their interactions. Though each analysis used only a subsets of imaging sessions, the representation of current trial events (Figure 5A,B top panels) was in both cases very similar to that for the full dataset (Figure 3E). As both the transition and reward probabilities determine which first step action is correct, effects common to these two analyses could in principle be mediated by changes in first-step action values rather than the reward or transition probabilities themselves, but effects that are specific to one or other analysis cannot.

**Figure 5.**
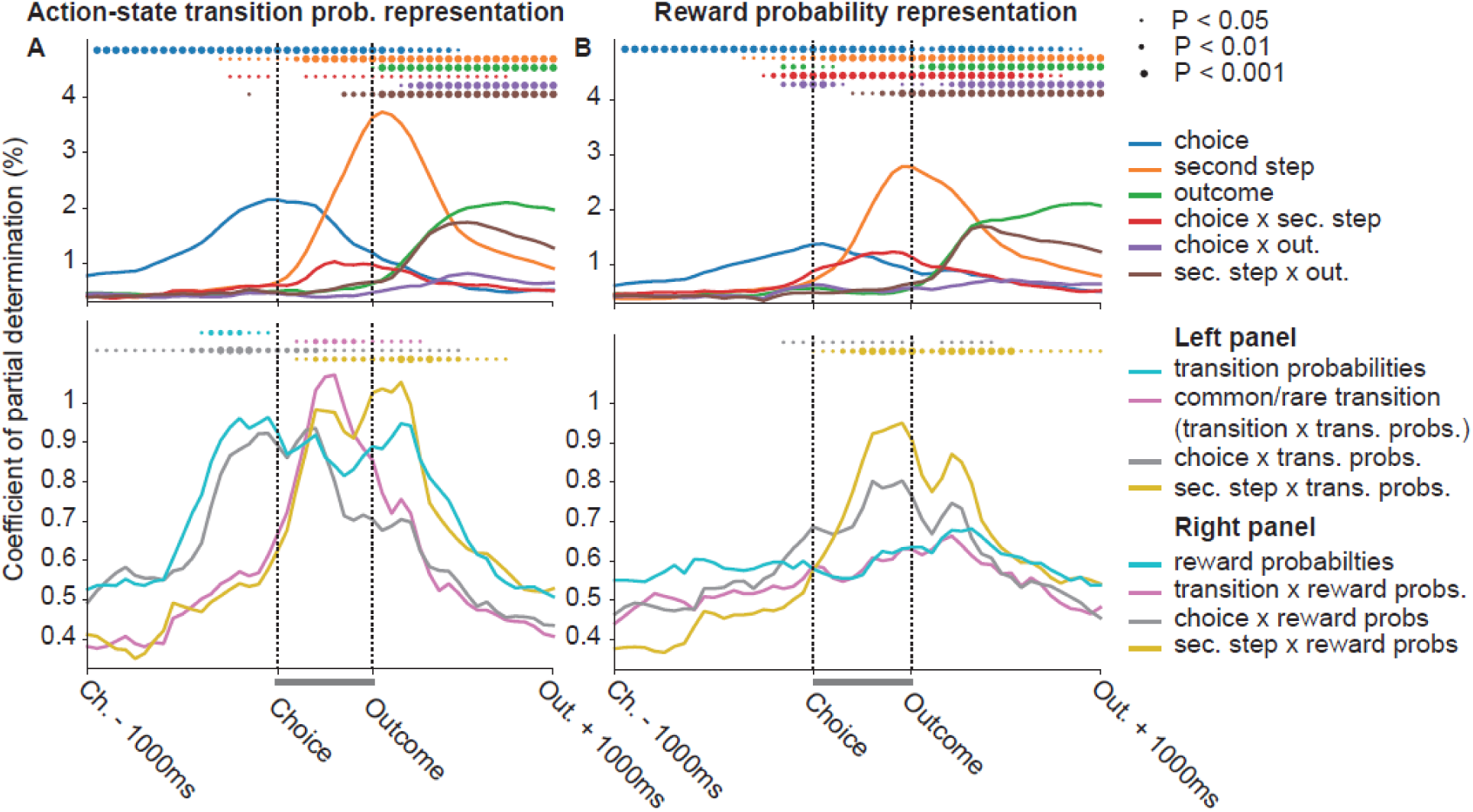
ACC represents model-based decision variables. **A**) Regression analysis predicting neuronal activity as a function of events on the current trial (top panel) and their interaction with the transition probabilities mapping the first-step choice to second-step states (bottom panel) for a subset of sessions with sufficient coverage of both states of the transition probabilities. Predictors plotted in top panels are as in figure 3E. Predictors plotted in the bottom panel are; *transition probabilities*: which of the two possible states the transition probabilities are in (see Fig. 1C), *common/rare transition*: whether the transition on the current trial was common or rare, i.e. the interaction of the transition on the current trial (e.g. top→right) with the state of the transition probabilities, *choice x trans. probs.*: the choice on the current trial interacted with the state of the transition probabilities – i.e. the predicted second-step state given the current choice, *sec. step x trans. probs.*: the second-step state reached on the current trial interacted with the state of the transition probabilities, i.e. the action which commonly leads to the second step state reached. Predictors shown in top and bottom panels of **A** were run as a single regression but plotted on separate axes for clarity. The grey bars between choice and outcome indicate the time period that was warped to align trials of different length. **B)** Regression analysis predicting neuronal activity as a function of events on the current trial (top panel) and their interaction with the reward probabilities in the second-step states (bottom panel) for a subset of sessions with sufficient coverage of different states of the reward probabilities. Predictors plotted in the bottom panel are; *reward probabilities*: which of the three possible states the transition probabilities are in (see Fig. 1C), *transition x reward probs*: Interaction of the transition on the current trial with the state of the reward probabilities. *choice x reward probs.*: the choice on the current trial interacted with the state of the reward probabilities, *sec. step x trans. probs.*: the second-step state reached on the current trial interacted with the state of the rewarded probabilities, i.e. the expected outcome (rewarded or not). Predictors shown in top and bottom panels of **B** were run as a single regression but plotted on separate axes for clarity.

Representation of the current state of the transition probabilities (Figure 5A: cyan), but not reward probabilities (Figure 5B: cyan), ramped up prior to choice and was sustained through trial outcome, though was only significant in the pre-choice period. Representation of the predicted second-step state given the current choice (the interaction of the choice on the current trial with the state of the transition probabilities) also ramped up prior to choice (Figure 5A: grey), peaking around choice time. Though ACC represented the interaction of choice with the reward probabilities (Figure 5B: grey), the time course was different, with weak representation prior to choice and a peak shortly before trial outcome.

Once the second-step state was revealed, ACC represented whether the transition was common or rare - i.e. the interaction of the transition on the current trial with the state of the transition probabilities (Figure 5A: magenta). There was no representation of the equivalent interaction of the transition on the current trial with the state of the reward probabilities (Figure 5B: magenta). Finally, ACC represented the interaction of the second-step state reached on the current trial with both the transition and reward probabilities, with both representations ramping up after the second-step state was revealed and persisting till after trial outcome (Figure 5 A,B: yellow). The interaction of second-step state with the transition probabilities corresponds to the action which commonly leads to the second-step state reached, potentially providing a substrate for model-based credit assignment. The interaction of second-step state with the reward probabilities corresponds to the predicted trial outcome (rewarded or not).

These data indicate that ACC represented a set of decision variables required for model-based RL, including the current action-state transition structure, the predicted state given chosen action, and whether the observed state transition was expected or surprising.

### Single-Trial Optogenetic Inhibition of Anterior Cingulate impairs model-based RL

To test the causal role of ACC in two-step task behaviour we silenced ACC neurons on individual trials using JAWS (Chuong et al., 2014). An AAV viral vector expressing JAWS-GFP under the CaMKII promotor was injected bilaterally into ACC of experimental animals (n = 11 mice, 192 sessions) (Figure S4), while GFP was expressed in control animals (n = 12 mice, 197 sessions). A red LED was chronically implanted above the cortical surface (Figure 6A). Electrophysiology confirmed that red light (50mW, 630nM) from the implanted LED robustly inhibited ACC neurons (Figure 6B, Kruskal-Wallis P < 0.05 for 67/249 recorded cells). ACC neurons were inhibited on a randomly selected 1/6 trials, with a minimum of two non-stimulated trials between each stimulation. Light was delivered from the time when the subject entered the side port and received the trial outcome until the time of the subsequent choice (Figure 6C).

**Figure 6.**
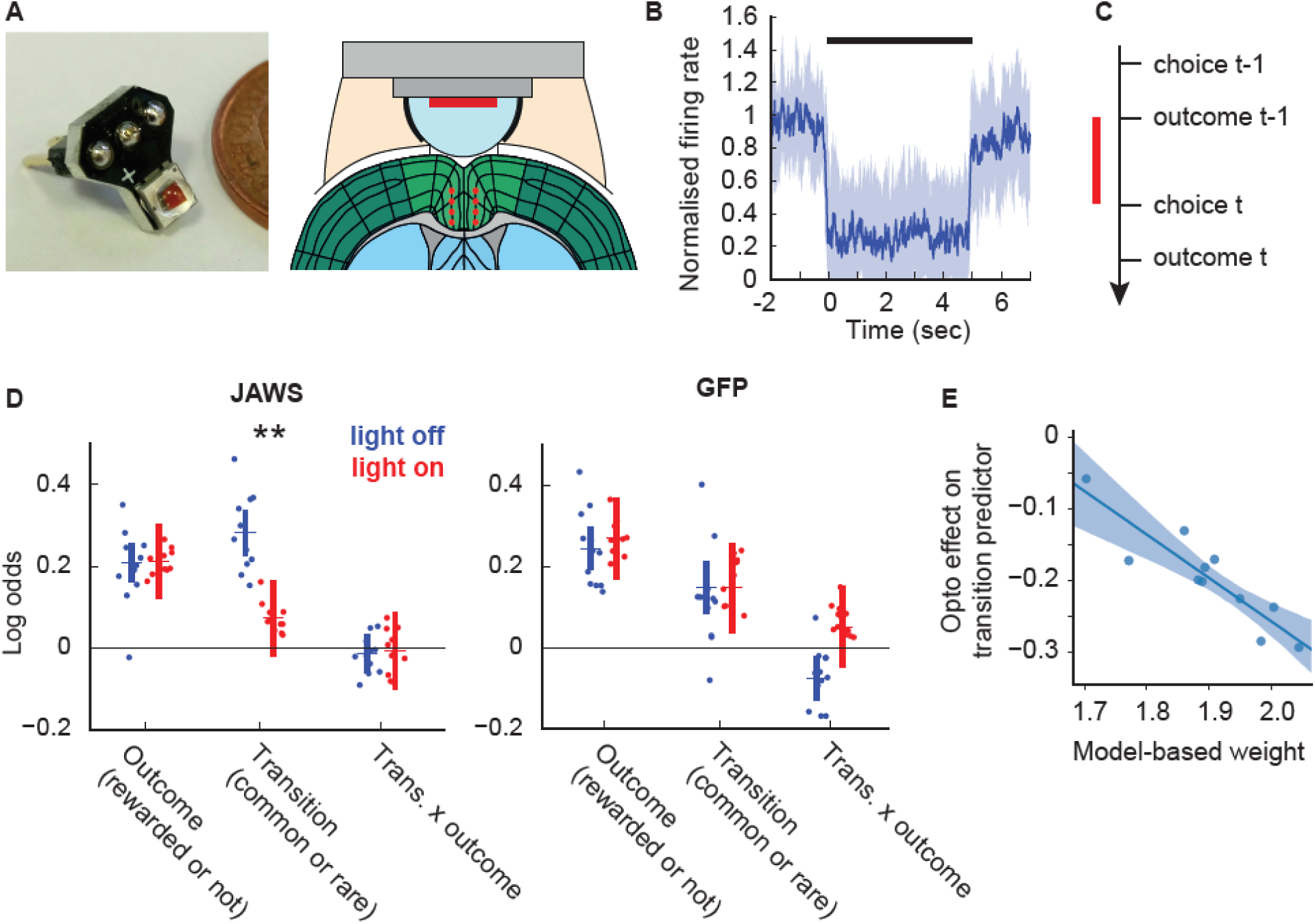
Optogenetic inhibition of ACC in the two-step task. **A)** LED implant (left) and diagram showing implant mounted on head (right), red dots on diagram indicate location of virus injections. **B)** Normalised firing rate for significantly inhibited cells over 5 second illumination, dark blue line – median, shaded area 25 – 75 percentiles **C)** Timing of stimulation relative to trial events. Stimulation was delivered from trial outcome to subsequent choice. **D)** Logistic regression analysis of ACC inhibition data showing loadings for the outcome, transition and transition-outcome interaction predictors for choices made on stimulated (red) and non-stimulated (blue) trials. ** indicates Bonferoni corrected P<0.01 between stimulated and non-stimulated trials. **E)** Correlation across subjects between the strength of model-based influence on choice (assessed using the RL model’s model-based weight parameter *G*_*mb*_) and the effect of optogenetic inhibition on the logistic regression model’s transition predictor.

ACC inhibition reduced the influence of the state transition (common or rare) on the subsequent choice (P = 0.007 Bonferroni corrected for comparison of 3 predictors, stimulation by group interaction P = 0.029, permutation test) (Figure 6D, S5A). Stimulation did not affect how either the trial outcome (P = 0.94 uncorrected), nor the transition-outcome interaction (P = 0.90 uncorrected) influenced the subsequent choice. In both experimental and control groups, light stimulation produced a bias towards the top poke, potentially reflecting an orienting response (bias predictor P < 0.001 uncorrected). Reaction times were not affected by light in either group (Paired t-test P > 0.36).

The selective impairment of the influence of action-state transition on subsequent choice, while sparing the influence of the trial outcome, is consistent with disrupted model-based control, as the transition predictor most strongly differentiates these two strategies (Figure 2). Consistent with this, the effect of inhibition on the transition predictor in each subject was strongly correlated with the strength of model-based influence on that subject’s choices (Figure 6E, R = -0.91, P = 0.0001), as assessed by fitting the RL model to subject’s behaviour in the inhibition sessions using a single set of parameters for all trials. Additional control analyses presented in supplementary results rule out an interpretation of the inhibition effect on the transition predictor in terms of motor level variables.

If ACC causally mediates model-based but not model-free RL, inhibiting ACC in a task where these strategies give similar recommendations should have little effect. To test this, we performed the same ACC manipulation in a probabilistic reversal learning task, where model-based and model-free RL are expected to generate qualitatively similar behaviour (supplementary results, Figure S6). ACC inhibition produced only a very subtle (but significant) reduction in the influence of the most recent outcome on the subsequent choice, suggesting that in this simpler task where model-based and model-free RL both recommend repeating rewarded choices, other regions could largely compensate for ACC inhibition.

## Discussion

We developed a novel two-step decision task for mice with reversals in the transition probabilities, designed to dissociate model-based and model free RL while rendering nugatory strategies based on latent-state inference. A detailed characterisation of subjects’ behaviour indicated that using this task we could quantify the usage of model-based and model-free RL in each subject. Calcium imaging indicated that different populations of ACC neurons represented each stage of the trial, with ramping choice selective activity followed by an evolving representation of the state reached and trial outcome. Representation of trial outcome (rewarded or not) was weaker than that of the state where the outcome was obtained, and different populations of neurons represented trial outcome in different states. ACC neurons represented a set of model-based decision variables, including the current action-state transition structure, the state predicted given the chosen action, and whether state transitions were expected or surprising. Consistent with this, optogenetic inhibition of ACC on individual trials reduced the influence of action-state transitions on subsequent choice, without affecting the influence of rewards. The strength of this inhibition effect strongly correlated across subjects with their use of model-based RL. These data demonstrate a role for ACC in model-based action selection.

Our study is one of several recent adaptations of two-step tasks for animal models (Miller et al., 2017; Dezfouli and Balleine, 2017; Hasz and Redish, 2018; Groman et al., 2019). Unlike these implementations, we introduced a major structural change to the task – reversals in the transition probabilities mapping first-step actions to second-step states. We did this to prevent subjects solving the task by inferring the current state of the reward probabilities (i.e. where rewards have recently been obtained) and learning fixed habitual strategies conditioned on this latent state (e.g. rewards on the left → choose up). We have previously shown that such strategies generate behaviour that looks very similar to model-based RL (Akam et al., 2015). This is a particular concern in animal two-step tasks. Human subjects are given detailed information about the structure of the task beforehand so they start with a largely correct model, then perform a limited number of trials with little contrast between good and bad options. Animal subjects are typically extensively trained, with strong contrast between good and bad options - giving ample opportunity and incentive to learn alternative strategies. In humans, extensive training renders apparently model-based behaviour resistant to a cognitive load manipulation (Economides et al., 2015) which normally disrupts model-based control (Otto et al., 2013), suggesting that it is possible to develop automatized strategies which closely resemble planning.

Introducing reversals in the transition probabilities breaks the long-term predictive relationship between where rewards are obtained and which first-step action has higher value. This precludes a habit-like strategy that exploits this simple relationship, but should not confound a model-based strategy beyond requiring ongoing learning about the current state of the transition probabilities. We compared behaviour on versions of the task with and without transition probability reversals, and found that this radically changed behaviour, both in terms of overall performance and the granular structure of learning. This strongly suggests that subjects used different strategies on the different versions, and while not conclusive, is consistent with the idea that with fixed transition probabilities, subjects learn sophisticated habits operating over the task’s latent state space. Another potentially confounding internal state description in this case is the successor representation (Dayan, 1993), which characterises current states in terms of their likely future. Successor representations support rapid updating of values in the face of changes in the reward function (and so could solve the fixed transition probability version of the task), but not changes in state transition probabilities (and so could not solve the new task) (Russek et al., 2017). Both of these strategies are of substantial interest in their own right, so understanding what underpins the behavioural differences between the task variants is a pressing question for future work.

It has been argued that differences in reaction time at the second-step following common vs rare transitions are additional evidence for model-based RL (Miller et al., 2017). However, in versions of the task where the actions required by different second-step states are consistent from trial to trial, reaction time differences may reflect preparatory activity at the level of the motor system, for example based on the strong correlation between the first-step choice and the *action* that will be required at the second-step. Indeed, a recent study using a two-step task in humans has shown that motor responses can show sensitivity to task structure even when choices are model-free (Konovalov and Krajbich, 2020). We therefore worry that second-step reaction times may not provide strong evidence that state prediction is used for model-based action evaluation.

As a starting point for neurophysiological investigation, we focused on a region of medial frontal cortex on the boundary between anterior-cingulate regions 24a and 24b and mid-cingulate regions 24a’ and 24b’ (Vogt and Paxinos, 2014). Though it has not to our knowledge been studied in the context of distinguishing actions and habits, there are anatomical physiological and lesion-based reasons in rodents, monkeys and humans for considering this particular role for the structure. First, neurons in rat (Sul et al., 2010) and monkey (Ito et al., 2003; Matsumoto et al., 2003; Kennerley et al., 2011; Cai and Padoa-Schioppa, 2012) ACC carry information about chosen actions, reward, action values and prediction errors during decision making tasks. Where reward type (juice flavour) and size were varied independently (Cai and Padoa-Schioppa, 2012), a subset of ACC neurons encoded the chosen reward type rather than the reward value, consistent with a role in learning action-state relationships. In a probabilistic decision making task in which reward probabilities changed in blocks, neuronal representations in rat ACC underwent abrupt changes when subjects detected a possible block transition (Karlsson et al., 2012). This suggests that the ACC may represent the block structure of the task, a form of world model used to guide action selection, albeit one based on learning about latent states of the world (Gershman and Niv, 2010; Akam et al., 2015), rather than the forward action-state transition model of classical model-based RL.

Second, neuroimaging in the original two-step task has identified representation of model-based value in anterior- and mid-cingulate regions, suggesting this is an important node in the model-based controller (Daw et al., 2011; Doll et al., 2015; Huang et al., 2020). Neuroimaging in a two-step task variant also found evidence for state prediction errors in dorsal ACC (Lockwood et al., 2019), consistent with our finding that ACC represented whether state transitions were common or rare. Relatedly, neuroimaging in a saccade task in which subjects constructed and updated a model of the location of target appearance found ACC activation when subjects updated an internal model of where saccade targets were likely to appear, (O’Reilly et al., 2013).

Third, ACC lesions in macaques produce deficits in tasks which require learning of action-outcome relationships (Hadland et al., 2003; Kennerley et al., 2006; Rudebeck et al., 2008), though the designs do not identify whether it is representation of the value or other dimensions of the outcome which were disrupted. Lesions of rodent ACC produce selective deficits in cost benefit decision making where subjects must weigh up effort against reward size (Walton et al., 2003; Rudebeck et al., 2006); however, again, the associative structures concerned are not clear.

Finally, the ACC provides a massive innervation to the posterior dorsomedial striatum (Oh et al., 2014; Hintiryan et al., 2016), a region necessary for learning and expression of goal directed action as assessed by outcome devaluation (Yin et al., 2005a, 2005b; Hilario et al., 2012).

Our study specifically tests the hypothesized role of ACC suggested by this body of work, by showing that ACC neurons represent variables critical for model-based RL, and that ACC activity is necessary for using action-state transitions to guide subsequent choice. More broadly, our study shows that it is possible to fashion sophisticated multi-step decision tasks that mice can acquire quickly and effectively, bringing to bear modern genetic tools to dissect mechanisms of model-based decision making.

## Acknowledgements

We thank Zach Mainen, Joe Patton, Mark Walton, Tim Behrens, Nathaniel Daw, Kevin Miller and Bruno Miranda for discussions about the work. The authors acknowledge the use of the Champalimaud Scientific and Technological Platforms and the University of Oxford Advanced Research Computing (ARC) facility (http://dx.doi.org/10.5281/zenodo.22558).

## Author contributions

Conceptualization: T.A., P.D., R.M.C., Investigation: T.A., I.R.V., I.M., X.Z., M.P., R.O., Data curation: T.A., I.M., M.P., R.O., Formal analysis: TA, Writing – original draft: T.A., Writing - review and editing T.A., P.D., R.M.C, Funding Acquisition: T.A., R.M.C, Supervision: P.D., R.M.C.

## Funding

TA was funded by the Wellcome Trust (WT096193AIA). RC was funded by the National Institute of Health (5U19NS104649) and ERC CoG (617142). PD was funded by the Gatsby Charitable Foundation, the Max Planck Society and the Humboldt Foundation. M.P., I.R.V. and I.M. were funded by the Fundação para a Ciência e Tecnologia (SFRH/BD/52222/2013, PD/BD/105950/2014, SFRH/BD/51715 /2011).

## Competing interests

The authors have no competing interests to report.

## Methods

### Experimental model and subject details

All procedures were reviewed and performed in accordance with the Champalimaud Centre for the Unknown Ethics Committee guidelines. 65 male C57BL mice aged between 2 – 3 months at the start of experiments were used in the study. Animals were housed under a 12 hours light/dark cycle with experiments performed during the light cycle. 17 subjects were used in the two-step task baseline behaviour dataset. 4 subjects were used in the ACC imaging. 2 subjects were used for electrophysiology controls for the optogenetics. 14 subjects (8 JAWS, 6 GFP controls) were used for the two-step task ACC manipulation only. 14 subjects (8 JAWS, 6 GFP controls) were used for the probabilistic reversal learning task ACC manipulation only. 14 subjects (8 JAWS, 6 GFP controls) were first trained and tested on the two-step ACC manipulation, then retrained for a week on the probabilistic reversal learning task and tested on the ACC manipulation in this task. 7 JAWS-GFP animals were excluded from the study due to poor or mis-located JAWS expression. In the group that was tested on both tasks, 1 Jaws and 2 control animals were lost from the study before optogenetic manipulation on the probabilistic reversal learning task due to failure of the LED implants. The resulting group sizes for the optogenetic manipulation experiments were as reported in the results section.

## Method details

### Behaviour

Mice were placed on water restriction 48 hours before the first behavioural training session, and given 1 hour ad libitum access to water in their home cage 24 hours before the first training session. Mice received 1 training session per day of duration 1.5 – 2 hours, and were trained 6 days per week with 1 hour *ad libitum* water access in their home cage on their day off. During behavioural training mice had access to dry chow in the testing apparatus as we found this increased the number of trials performed and amount of water consumed. On days when mice were trained they typically received all their water in the task (typically 0.5-1.25ml), but additional water was provided as required to maintain a body weight >85% of their pre-restriction weight. Under this protocol, bodyweight typically dropped to ∼90% of pre-restriction level in the first week of training, then gradually increased over weeks to reach a steady state of ∼95-105% pre-restriction body weight.

Behavioural experiments were performed in 14 custom made 12×12cm operant chambers using pyControl (http://pycontrol.readthedocs.io/), a behavioural experiment control system built around the Micropython microcontroller.

### Two-step task

The apparatus, trial structure and block structure of the two-step task are described in the results section. Block transitions were triggered based on subject’s behaviour, occurring 20 trials after an exponential moving average (tau = 8 trials) of subject’s choices crossed a 75% correct threshold. The 20 trial delay between the threshold crossing and block transition allowed subjects performance at the end of blocks to be assessed without selection bias due to the block transition rule. In neutral blocks where there was no correct choice, block transitions occurred with 0.1 probability on each trial after the 40^th^, giving a mean neutral block length of 50 trials. Subjects started each session with the reward and transition probabilities in the same state that the previous session finished on.

Subjects encountered the full trial structure from the first day of training. The only task parameters that were changed over the course of training were the reward and state transition probabilities and the reward sizes. These were changed to gradually increase task difficulty over days of training, with this typical trajectory of parameter changes shown in table 1.

**Table 1:** Two-step task parameter changes over training.

| Table 1: Two-step task parameter changes over training |  |  |  |
| --- | --- | --- | --- |
| Session number | Reward size (ul) | Transition probabilities (common / rare) | Reward probabilities (good / bad side) |
| 1 | 10 | 0.9 / 0.1 | First 40 trials all rewarded, subsequently 0.9 / 0.1 |
| 2 - 4 | 10 | 0.9 / 0.1 | 0.9 / 0.1 |
| 5 - 6 | 6.5 | 0.9 / 0.1 | 0.9 / 0.1 |
| 7 - 8 | 4 | 0.9 / 0.1 | 0.9 / 0.1 |
| 9 - 12 | 4 | 0.8 / 0.2 | 0.9 / 0.1 |
| 13+ | 4 | 0.8 / 0.2 | 0.8 / 0.2 |

### Probabilistic reversal learning task

Mice were trained to initiate each trial in a central nose-poke port which was flanked by left and right poke ports. Trial initiation caused the left and right pokes to light up and subjects then chose between them for the chance of obtaining a water reward. Reward probabilities changed in blocks, with three block types; *left good* (left=0.75/right=0.25), *neutral* (0.5/0.5) and *right good* (0.25/0.75). Block transitions from non-neutral blocks were triggered 10 trials after an exponential moving average (tau = 8 trials) crossed a 75% correct threshold. Block transitions from neutral blocks occurred with probability 0.1 on each trial after the 15^th^ of the block to give an average neutral block length of 25 trials.

#### Optogenetic Inhibition

Experimental animals were injected bilaterally with *AAV5-CamKII-Jaws-KGC-GFP-ER2* (UNC vector core, titre: 5.9 x 10^12^) using 16 injections each of 50nL (total 800nL) spread across 4 injection tracks (2 per hemisphere) at coordinates: AP: 0, 0.5, ML: ±0.4, DV: -1, -1.2, -1.4, -1.6mm relative to dura. Control animals were injected with *AAV5-CaMKII-GFP* (UNC vector core, titre: 2.9 x 10^12^) at the same coordinates. Injections were performed at a rate of 4.6nL/5 seconds, using a Nanojet II (Drummond Scientific) with bevelled glass micropipettes of tip diameter 50-100um. A circular craniotomy of diameter 1.8mm was centred on AP: 0.25, ML: 0, and a high power red led (Cree XLamp XP-E2) was positioned above the craniotomy touching the dura. The LED was mounted on a custom designed insulated metal substrate PCB (Figure 6A). The LEDs were powered using a custom designed constant current LED driver. Light stimulation (50mW, 630nM) was delivered on stimulation trials from when the subject entered the side poke until the subsequent choice, up to a maximum of 6 seconds. Stimulation was delivered on a randomly selected 17% of trials, with a minimum of 2 non-stimulated trials between each stimulation trial followed by a 0.25 probability of stimulation on each subsequent trial. At the end of behavioural experiments, animals were sacrificed and perfused with paraformaldehyde (4%). The brains were sectioned in 50um coronal slices and the location of viral expression was characterised with fluorescence microscopy (Figure S4).

Two animals were injected unilaterally with the JAWS-GFP virus using the coordinates described above and implanted with the LED implant and a movable bundle of 16 tungsten micro-wires of 23μm diameter (Innovative-Neurophysiology) to record unit activity. After 4 weeks of recovery, recording sessions were performed at 24 hour intervals and the electrode bundle was advanced by 50 um after each session, covering a depth range of 300 – 1300um from dura over the course of recordings. During recording sessions mice were free to move inside a sound attenuating chamber. Light pulses (50mW power, 5 second duration) were delivered at random intervals with a mean inter-stimulus interval of 30 seconds. Neural activity was acquired using a Plexon recording system running Omniplex v. 1.11.3. The signals were digitally recorded at 40000 Hz and subsequently band-pass filtered between 200 Hz and 3000 Hz. Following filtering, spikes were detected using an amplitude threshold set at twice the standard deviation of the bandpass filtered signal. Initial sorting was performed automatically using Kilosort (Pachitariu et al., 2016). The results were refined via manual sorting based on waveform characteristics, PCA and inter-spike interval histogram. Clusters were classified as single units if well separated from noise and other units and the spike rate in the 2ms following each spike was less than 1% of the average spike rate.

#### ACC imaging

Mice were anaesthetized with a mix of 1-1.5% isofluorane and oxygen (1 l.min-1), while body temperature was monitored and maintained at 33°C using a temperature controller (ATC1000, World Precision Instruments). Unilateral injection of 300 nl of AAV5.αCaMKII.GCaMP6f.WPRE.SV40 (titer: 2.43×10^13^, Penn Vector Core) into the right Anterior Cingulate Cortex (AP: +1.0 mm; ML: +0.45mm; DV: -1.4mm) was performed using a Nanojet II Injector (Drummond Scientific, USA) at a rate of 4.6 nl per pulse, every 5 s. Injection pipette was left in place 20 min post-injection before removal. 25 minutes after injection, a 1mm diameter circular craniotomy was centered at coordinates (AP: +1.0 mm; ML: +0.55mm) and a 1mm GRIN lens (Inscopix) was implanted above the injection site at a depth of -1.2 mm ventral to the surface, and secured to the skull using cyanoacrylate (Loctite) and black dental cement (Ortho-Jet, Lang Dental USA). One 1/16-inch stainless-steel screw (Antrin miniatures) was attached to the skull to secure the cement cap that fixed the lens to the skull. Mice were then given an i.p. injection of buprenorfin (Bupaq, 0.1 mg.kg-1) and allowed to recover from anaesthesia in a heating mat before returning to home cage.

Three to four weeks after surgery, mice were anaesthetized and placed in the stereotactic frame, where a miniaturized fluorescence microscope (Inscopix) attached to a magnetic baseplate (Inscopix) were lowered to the top of the implanted GRIN lens, until a sharp image of anatomical landmarks (blood vessels) and putative neurons appeared in the focal plane. Baseplate was then cemented to the original head cap, allowing to fix the set focal plane for imaging.

For image acquisition during task behaviour, mice were briefly anaesthetized using a mixture of isofluorane (0.5-1%) and oxygen (1 l.min-1) and the miniaturized microscope was attached and secured to the baseplate. This was followed by a 20-30 min period of recovery in the home cage before imaging experiments. Image acquisition (nVistaHD, Inscopix) was done at 10 Hz, with LED power set to 10-30% (0.1-0.3 mW) with a gain of 3. Image acquisition parameters were set to the same values between sessions for each mouse.

### Quantification and statistical analysis

All analysis of behavioural data was performed in Python 3.

#### Logistic regression

Binary predictors used in logistic regressions are shown in table 2. The two-step task previous trial logistic regression (Figure 2B) used all predictors in table 1. The two-step task lagged logistic regression used predictors *Choice, Outcome, Transition* and *Transition-outcome interaction* at lags 1, 2, 3-4, 5-8, 8-12 (where lag 3-4 etc. means the sum of the individual trial predictors over the specified range of lags) and predictors *Bias: top/bottom*, and *Bias:clockwise/counter-clockwise*. The *Correct* predictors was included in the previous trial regression to prevent correlations across trials from causing spurious loading on the *Transition-outcome interaction* predictor (see Akam et. al. 2015 for discussion). It was not included in the lagged regression as here the effect of earlier trials is accounted for by the lagged predictors. For the two-step task regressions, the first 20 trials after each reversal in the transition probabilities was exclude for the analysis as it is ambiguous which transitions are common and rare at this point. This resulted in ∼9% of trials being excluded.

**Table 2:**
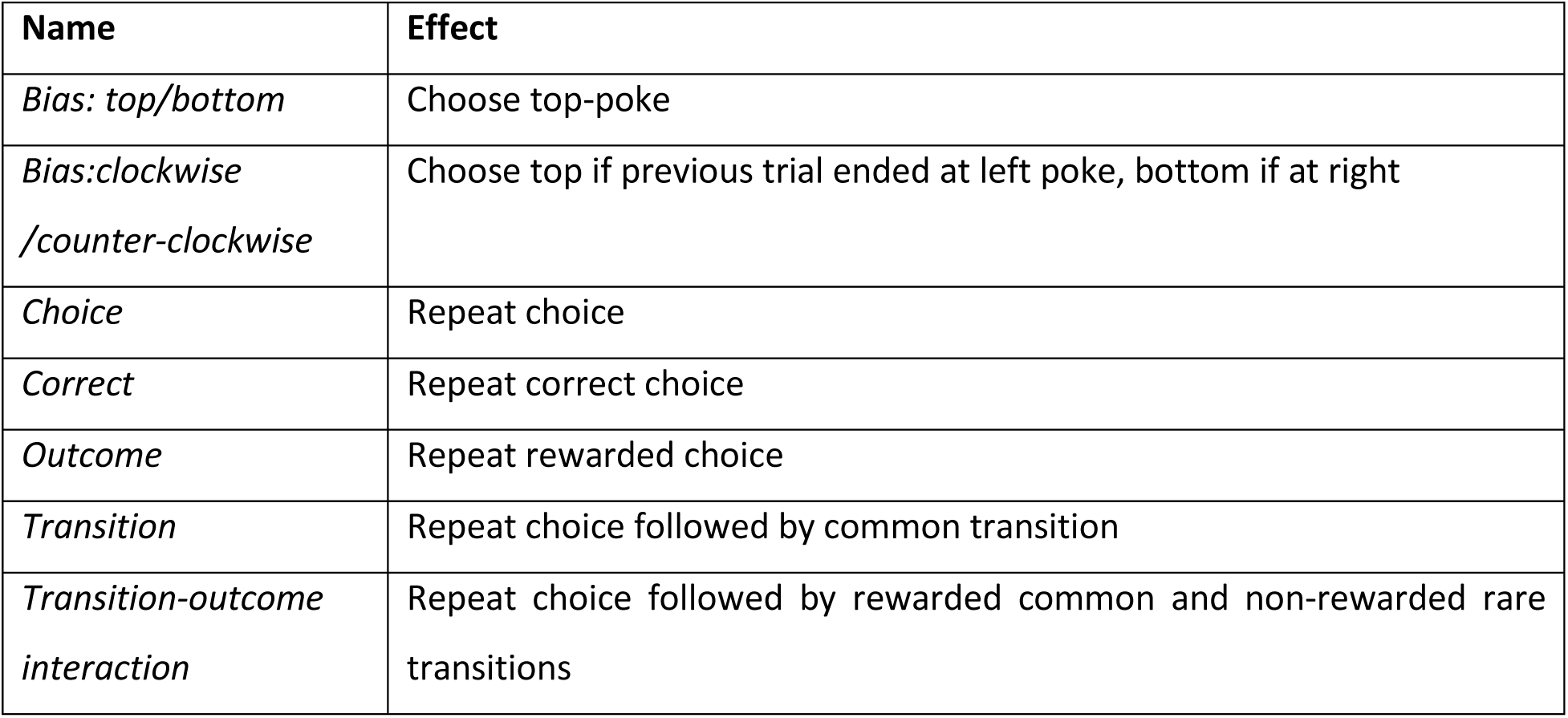
Predictors used in two-step task logistic regression.

| Table 2: Predictors used in two-step task logistic regression |  |
| --- | --- |
| Name | Effect |
| *Bias: top/bottom* | Choose top-poke |
| *Bias:clockwise /counter-clockwise* | Choose top if previous trial ended at left poke, bottom if at right |
| *Choice* | Repeat choice |
| *Correct* | Repeat correct choice |
| *Outcome* | Repeat rewarded choice |
| *Transition* | Repeat choice followed by common transition |
| *Transition-outcome interaction* | Repeat choice followed by rewarded common and non-rewarded rare transitions |

The logistic regression analysis for the probabilistic reversal learning task (Figure S6D) used predictors *Choice*, and *Outcome* at lags 1, 2, 3.

#### Reinforcement learning models

RL model variables and parameters are listed in table 3.

**Table 3:**
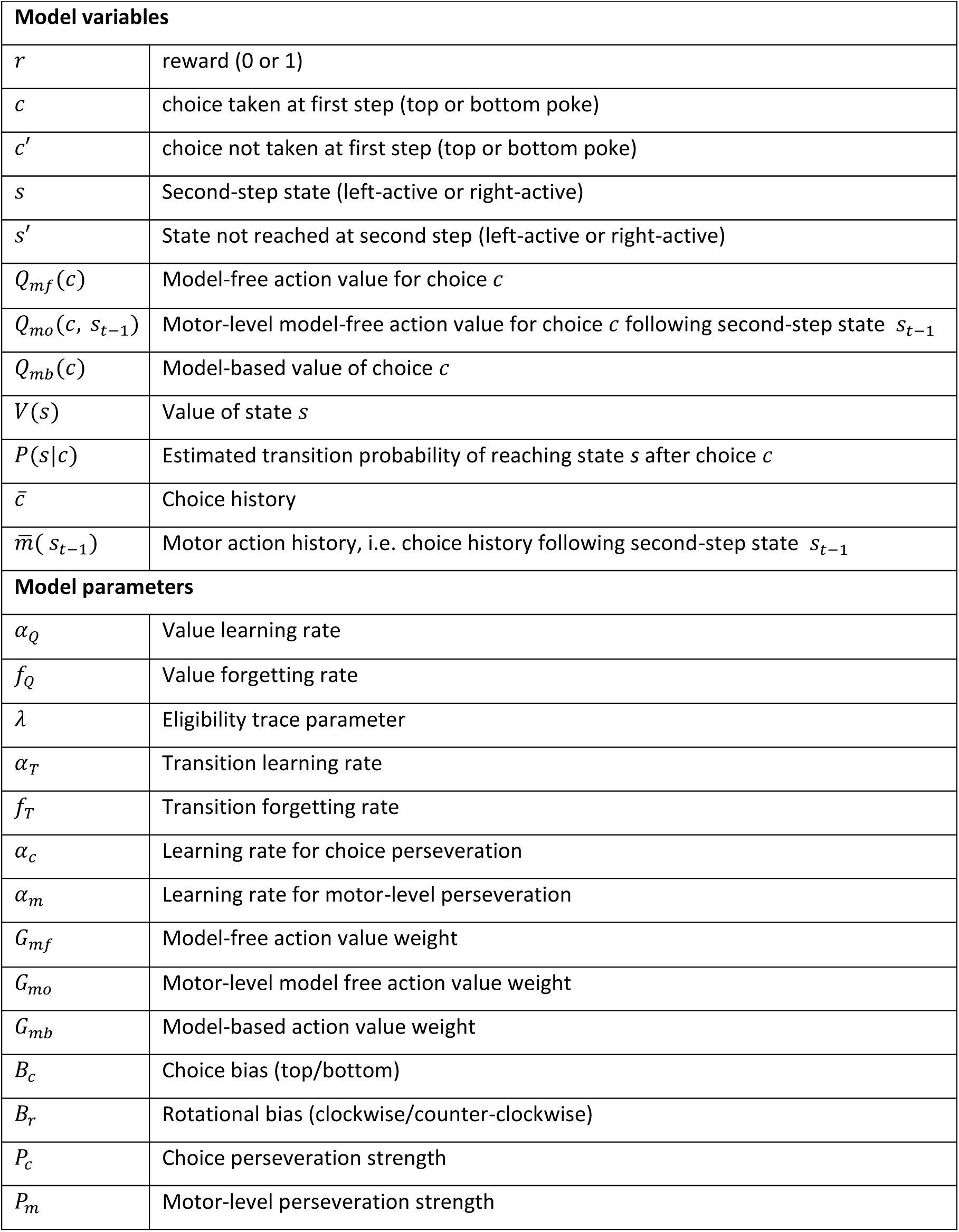
RL model variables and parameters.

Choice and state values were updated as:

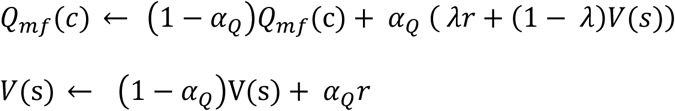

In models that included value forgetting this was implemented as:

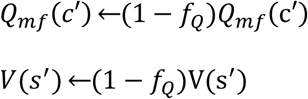

Action-state transition probabilities used by the model-based system were updated as:

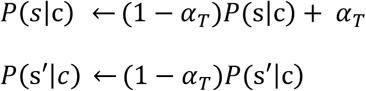

In models that included transition probability forgetting this was implemented as:

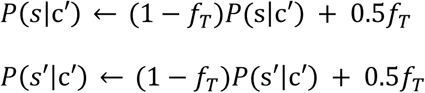

At the start of each trial, model-based first step action values were calculated as:

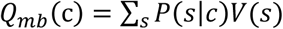

Models that included model-free values for first step motor actions (e.g. left→top), updated these as:

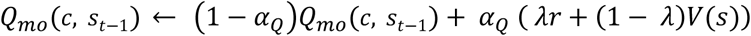

Motor level model-free value forgetting was implemented as:

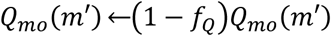

Where *m′* are all motor actions not taken.

Choice perseveration was modelled using a choice history variable 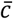. In models using single trial perseveration this was:

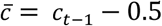

where *c*_*t*−1_ = 1 if previous choice is top and 0 if previous choice is bottom.

In models using multi-trial perseveration 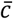 was an exponential moving average of recent choices, updated as:

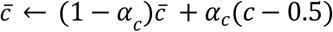

where *c* = 1 if choice is top and *c* = 0 if choice is bottom.

In models which used motor-level perseveration this was modelled using variables 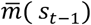 which were exponential moving averages of choices following trials ending in state *s*_*t*−1_, updated as:

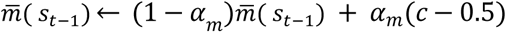

Net action values were given by a weighted sum of model-free, motor-level model-free and model-based action values, biases and perseveration.

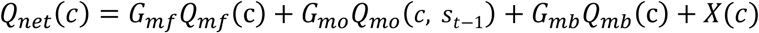

Where *G*_*mf*_, *G*_*mo*_ and *G*_*mb*_ are weights controlling the influence of respectively the model-free, motor-level model-free and model-based action values, and *X*(*c*) is biases and perseveration where:

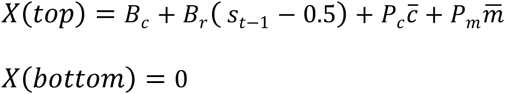

where *s*_*t*−1_ = 1 if previous second step state is left and 0 if right.

Net action values determined choice probabilities via the softmax decision rule:

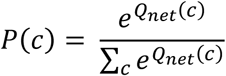

#### Hierarchical modelling

Both the logistic regression analyses and reinforcement learning model fitting used a Bayesian hierarchical modelling framework (Huys et al., 2011), in which parameter vectors ***h***_*i*_ for individual sessions were assumed to be drawn from Gaussian distributions at the population level with means and variance ***θ*** = **{*μ, Σ*}**. The population level prior distributions were set to their maximum likelihood estimate:

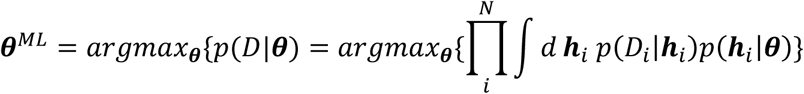

Optimisation was performed using the Expectation-Maximisation algorithm with a Laplace approximation for the E-step at the k-th iteration given by:

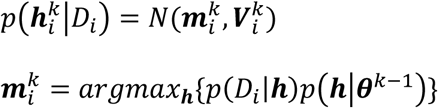

Where 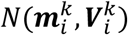 is a normal distribution with mean 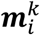 given by the maximum a posteriori value of the session parameter vector ***h***_*i*_ given the population level means and variance ***θ***^*k*−1^, and the covariance 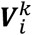 given by the inverse Hessian of the likelihood around 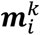. For simplicity we assumed that the population level covariance ***Σ*** had zero off-diagonal terms. For the k-th M-step of the EM algorithm the population level prior distribution parameters ***θ*** = **{*μ, Σ*}** are updated as:

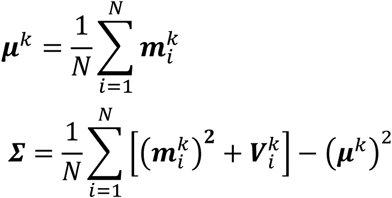

Parameters were transformed before inference to enforce constraints (0 < **{** *G*_*mf*_, *G*_*mo*_, *G*_*mb*_**}**, 0 < **{** *α*_*Q*_, *f*_*Q*_, *λ, α*_*T*_, *f*_*T*_, *α*_*c*_, *α*_*m*_**}** < 1).

#### Model comparison

To compare the goodness of fit for models with different numbers of parameters we used the integrated Bayes Information Criterion (iBIC) score. The iBIC score is related to the model log likelihood *p*(*D|M*) as:

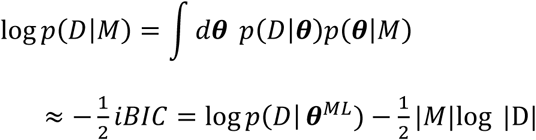

Where |M| is the number of fitted parameters of the prior, |D| is the number of data points (total choices made by all subjects) and iBIC is the integrated BIC score. The log data likelihood given maximum likelihood parameters for the prior *log p*(*D|* ***θ***^*ML*^) is calculated by integrating out the individual session parameters:

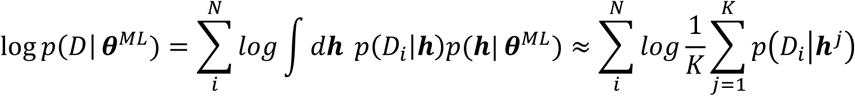

Where the integral is approximated as the average over K samples drawn from the prior *p*(***h****|****θ***^*ML*^). Bootstrap 95% confidence intervals were estimated for the iBIC scores by resampling from the population of samples drawn from the prior.

#### Permutation testing

Permutation testing was used to assess the significance of differences in model fits between stimulated and non-stimulated trials. The regression model was fit separately to stimulated and non-stimulated trials to give two sets of population level parameters ***θ***_*s*_ = **{*μ***_*s*_, ***Σ***_*s*_**}** and ***θ***_*n*_ = **{*μ***_*n*_, ***Σ***_*n*_**}**, where ***θ***_*s*_ are the parameters for the stimulated trials and ***θ***_*n*_ are the parameters for the non-stimulated trials. The difference between the population level means for the stimulated and non-stimulated conditions were calculated as:

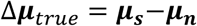

An ensemble of *N* = *5*000 permuted datasets was then created by shuffling the labels on trials such that trials were randomly assigned to the ‘stimulated’ and ‘non-stimulated’ conditions. The model was fit separately to the stimulated and non-stimulated trials for each permuted dataset and the difference between population level means in the stimulated and non-stimulated conditions was calculated for each permuted dataset *i* as:

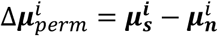

The distribution of Δ***μ***_*perm*_ over the population of permuted datasets approximates the distribution under the null hypothesis that stimulation does not affect the model parameters. The P-values for the observed distances Δ***μ***_*true*_ are then given by:

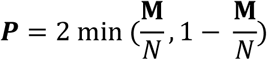

Where **M** is the number of permutations for which 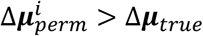.

In addition to testing for a significant main effect of the stimulation we tested for significant stimulation by group interaction. We first evaluated the true difference between the effect sizes for the two groups as:

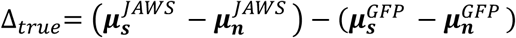

The approximate distribution of this difference under the null hypothesis that there was no difference between the groups was evaluated by creating an ensemble of permuted datasets in which we randomly assigned subjects to the JAWS and GFP groups and the interaction P value was calculated as above.

Permutation testing was also used to assess significance differences in logistic regression model fits to the behaviour of subjects run on the task variants with and without reversals in the transition probability reversals, with permuted datasets generated by permuting subjects between the two groups.

#### Bootstrap tests

To test whether logistic regression predictor loadings were significantly different from zero, bootstrap confidence intervals on the population means ***μ*** were evaluated by generating a set of *N* = 5000 resampled datasets by sampling subjects with replacement. P values for predictor loading significantly different from zero were calculated as:

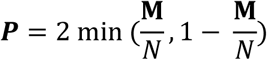

Where **M** is the number of resampled datasets for which ***μ***>0.

#### Analysis of simulated data

For analyses of data simulated from different RL agent types (Figure 2), we first fitted each agent to our baseline behavioural dataset using the hierarchical framework outlined above. The agents used were a model-free agent with eligibility traces and value forgetting (Figure 2D-F), and a model-based agent with value and transition probability forgetting (Figure 2G-I) and the best fitting RL model described in supplementary results (Figure 2J-L). We then simulated data (4000 sessions each of 500 trials) from each agent, drawing parameters for each session from the fitted population level distributions for that agent. We performed the logistic regression on the simulated data, using the same hierarchical framework as for the experimental data.

#### Calcium imaging analysis

##### Pre-processing

All imaging videos were pre-processed and motion corrected using custom MATLAB code, using the Mosaic API (Inscopix). Videos were spatially down sampled 4×4 and motion corrected using a 15 to 20-point specific reference area drawn for each animal (blood vessel pattern). Black pixel borders inserted during motion correction were then removed by cropping the corrected videos.

To extract calcium signals from putative single neurons, we used the MATLAB implementation of the Constrained non-negative matrix factorization – extended algorithm (CNMF-E) (Zhou et al., 2018). Putative single units were isolated from the processed imaging videos and subsequently inspected manually for quality assessment of both spatial masks and calcium time series. Isolated putative units not matching spatial masks or temporal features of neurons were discarded and not used in following analyses. All analyses used the deconvolved activity inferred by CNMF-E. For the regression and trajectory analyses the deconvolved activity was log2 transformed. Activity was aligned across trials by warping the time period between the choice and second-step port entry to match the median trial timings, activity prior to choice and after second-step port entry was not warped. Following time warping, activity was up-sampled to 20Hz and Gaussian smoothed with 50ms standard deviation. Example activity before and after alignment and smoothing are shown in figure S7.

##### Regression analysis of neuronal activity

The regression analysis in figure 3E-H used binary predictors coding the choice (top or bottom), second-step state (right or left) and trial outcome (rewarded or not), as well as the two-way interactions of these predictors (e.g. choice x second-step). To assess whether coefficients of partial determination were significantly different from that expected by chance, we generated an ensemble of 5000 permuted datasets by circularly shifting the predictors relative to the neural activity by a random number of trials drawn independently for each session from the range [0, N] where N is the number of trials in the session. This permutation preserves the autocorrelation across trials in both the neural activity and the predictors but randomises the relationship between them. We calculated P values for each predictor at each time point as the fraction of permutations for which the permuted datasets had a larger CPD than the true dataset. P values for each predictor were corrected for multiple comparison across time-points using the Benjamini–Hochberg procedure (Benjamini and Hochberg, 1995).

In figure 3G we evaluated the time course for two orthogonal representations of second-step state which occurred pre- and post-trial outcome. We defined unit projection vectors from the regression weights for second-step state at a time point mid-way between choice and outcome and 250ms after outcome. We then projected the regression weights for second-step state at each time point onto these two vectors to obtain time-courses for each representation. To avoid selection bias distorting the time-courses, we divided the data into odd and even trials and used the odd trials to define projection vectors that weights from the even trials were projected onto, and vice versa.

In Figure 5A we used an additional binary predictor coding the state of the transition probabilities (*top→ right / bottom→ left* vs *top→ left / bottom→ right*), binary predictors coding the interaction of the transition probabilities with the choice and second step, and the transition on the current trial coded clockwise (e.g. top→right) vs counter-clockwise – i.e. whether the transition was common or rare. In figure 5B we used a predictor which coded the state of the reward probabilities as -0.5, 0, 0.5 for the *left-good, neutral* and *right-good* states respectively, as well as the interactions of this predictor with the choice, second-step and transition on the current trial. As the subjects knowledge of the transition/reward probabilities is ambiguous in the period following block transitions where they change, these predictors were coded 0 in the 20 trials following such changes, and ±0.5 at other times. These analyses included only sessions where we had at least 40 trials in at least two different states of the transition (Figure 5A) or reward (Figure 5B) probabilities.

##### Neuronal trajectory analysis

The activity trajectories in figure 4 were obtained by projecting the average population activity for each trial type into the low dimensional space that captured most variance between trial types, where trial type was defined by the 8 possible combinations of choice, second-step and outcome. To find this space, we calculated the average activity for each neuron for each trial type. We then averaged these across trial types to evaluate the component of activity that was not selective to different trial types. We subtracted the non-selective activity for each neuron from that neurons average activity for each individual trial type, and concatenated across trial types to generate a data matrix of shape [n neurons, n trial types * n time point] representing how activity for each neuron deviated from its cross-trial-type average in each trial type. We performed PCA on this matrix to find the space that captured the most cross-trial-type variance and then projected the average population activity trajectory for each trial type into this space to generate figure 4.

## Supplementary figures

**Figure S1.**
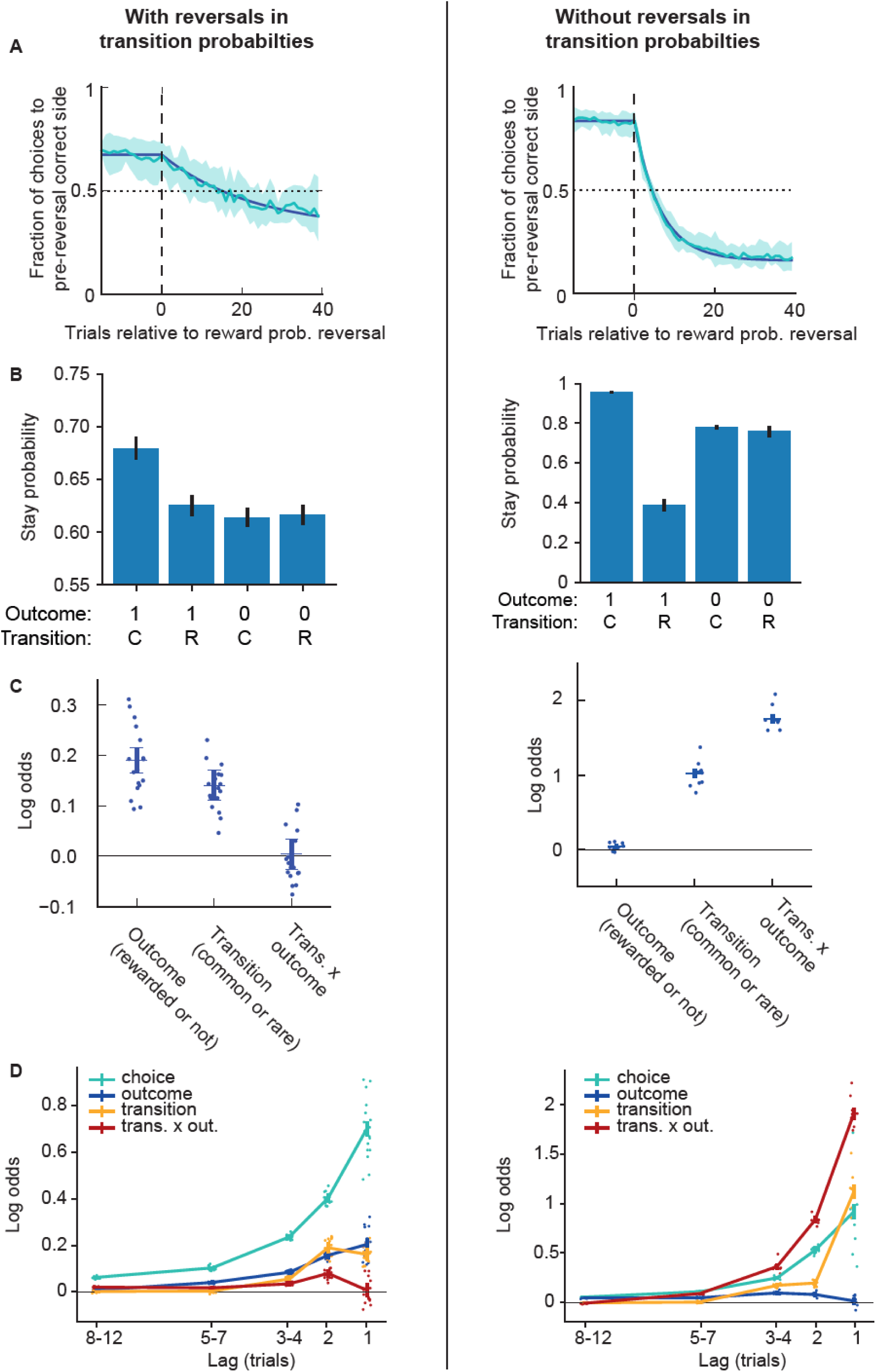
Behaviour without transition probability reversals. Comparison of behaviour a version of the two-step task with transition probability reversals (left panels – reproduced from figures 1 and 2 for ease of comparison) and without transition probability reversals (right panel). The tasks were identical apart from the presence/absence of transition probability reversals. **A)** Choice probability trajectories around reward probability reversals. Pale blue line – average trajectory, dark blue line – exponential fit, shaded area – cross-subject standard deviation. **B)** Stay probability analysis showing the fraction of trials the subject repeated the same choice following each combination of trial outcome (rewarded (1) or not (0)) and transition (common (C) or rare (R)). Error bars show cross-subject SEM. **C)** Logistic regression model fit predicting choice as a function of the previous trial’s events. Predictor loadings plotted are; *outcome* (repeat choices following rewards), *transition* (repeat choices following common transitions) and *transition-outcome interaction* (repeat choices following rewarded common transition trials and non-rewarded rare transition trials). Error bars indicate 95% confidence intervals on the population mean, dots indicate maximum a posteriori (MAP) subject fits. **D)** Lagged logistic regression model predicting choice as a function of events over the previous 12 trials. Predictors are as in **C**, predictor loading at lag *x* indicates the influence of events at trial *t* on choice at trial *t* + *x.*

**Figure S2.**
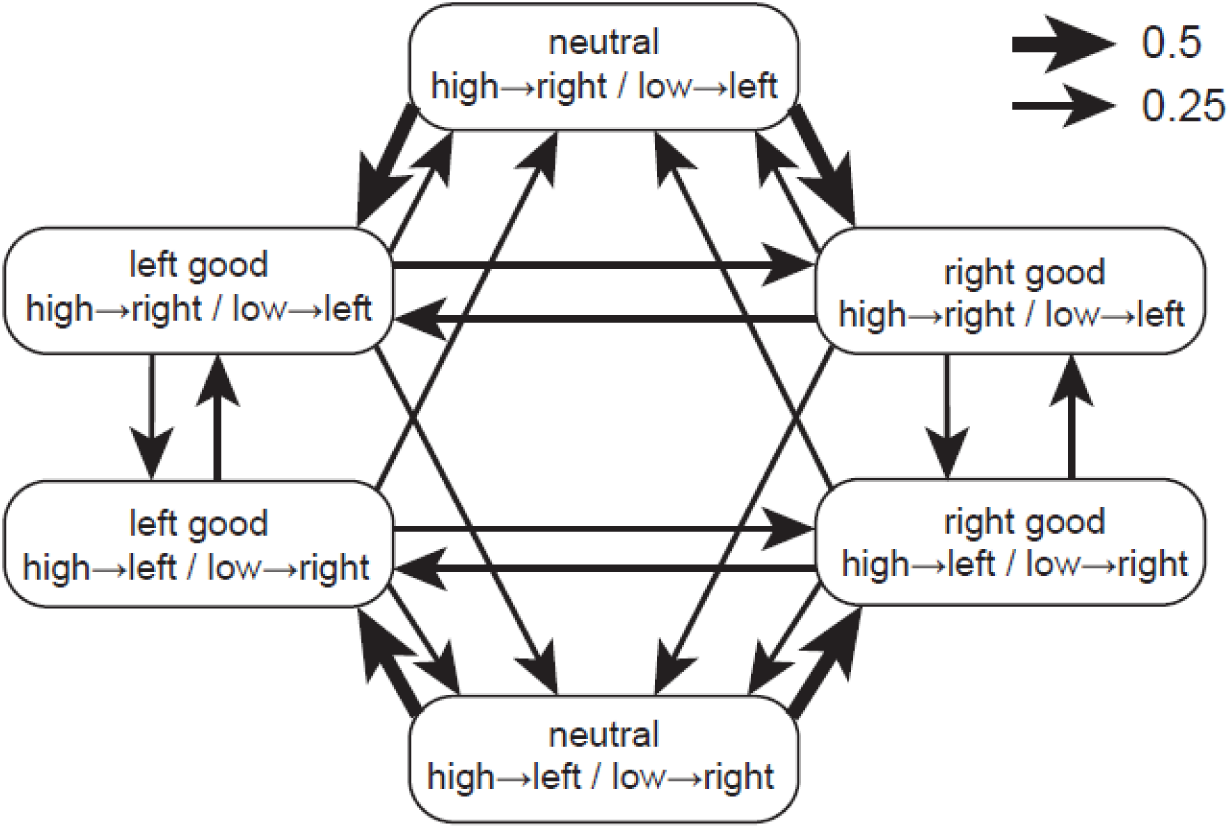
Block transition probabilities. Diagram of block transition probabilities for the two-step task.

**Figure S3.**
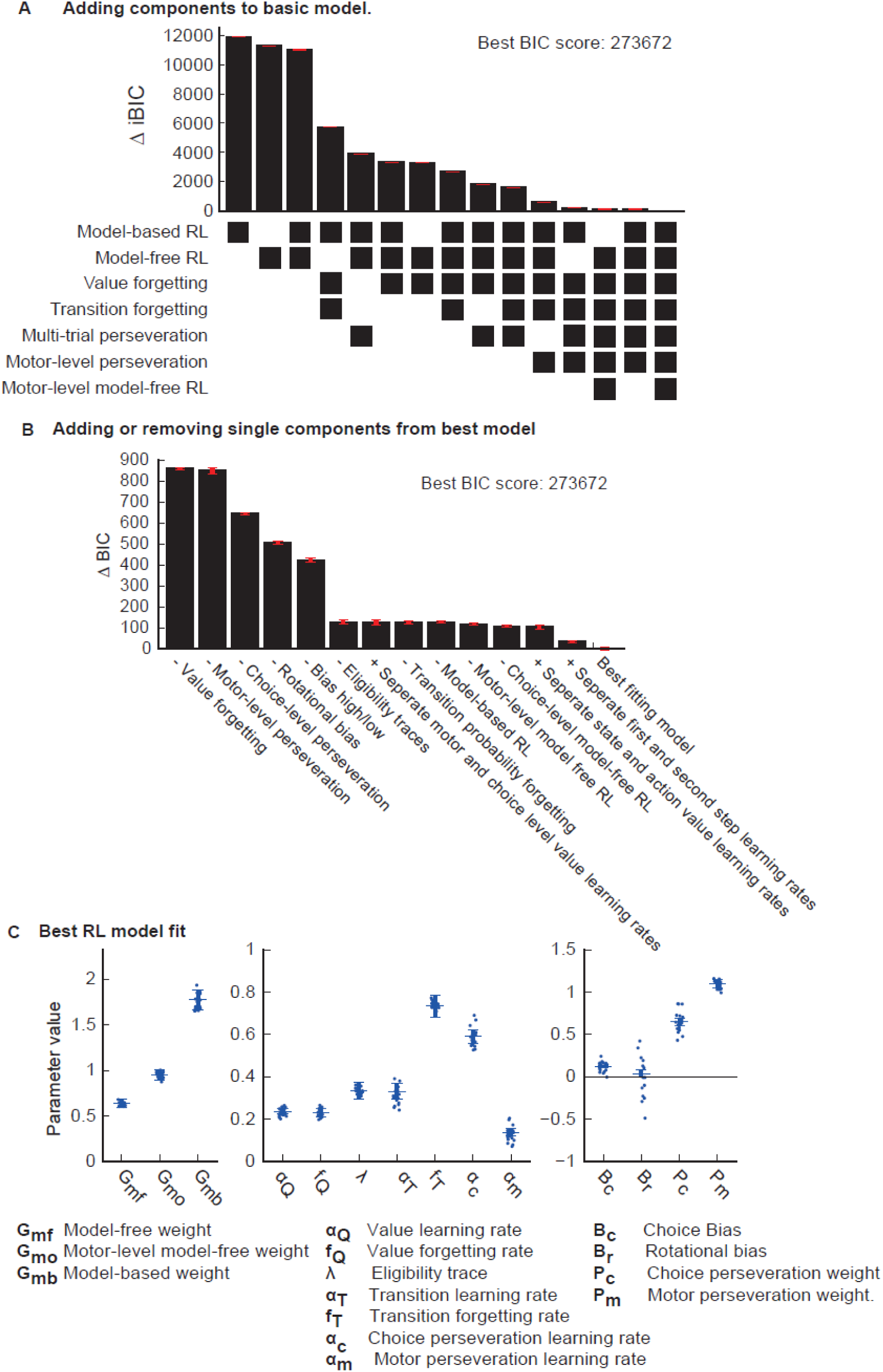
Baseline dataset BIC score model comparison. **A)** iBIC score comparison for set of RL models on baseline behavioural dataset. The set of models was constructed as described in supplementary results by iteratively adding features to the RL model. The grid below the plot indicates which features were included in each model. **B)** iBIC score comparison on the baseline dataset for set of RL models created by adding or removing a single feature at a time from the best fitting model. The text below each bar indicates what feature has been added or removed. Error-bars indicate the bootstrap 95% confidence interval on the BIC score. **C)** Parameter values for best fitting RL model. Bars indicate 95% confidence intervals on the population mean, dots indicate maximum a posteriori (MAP) subject fits.

**Figure S4.**
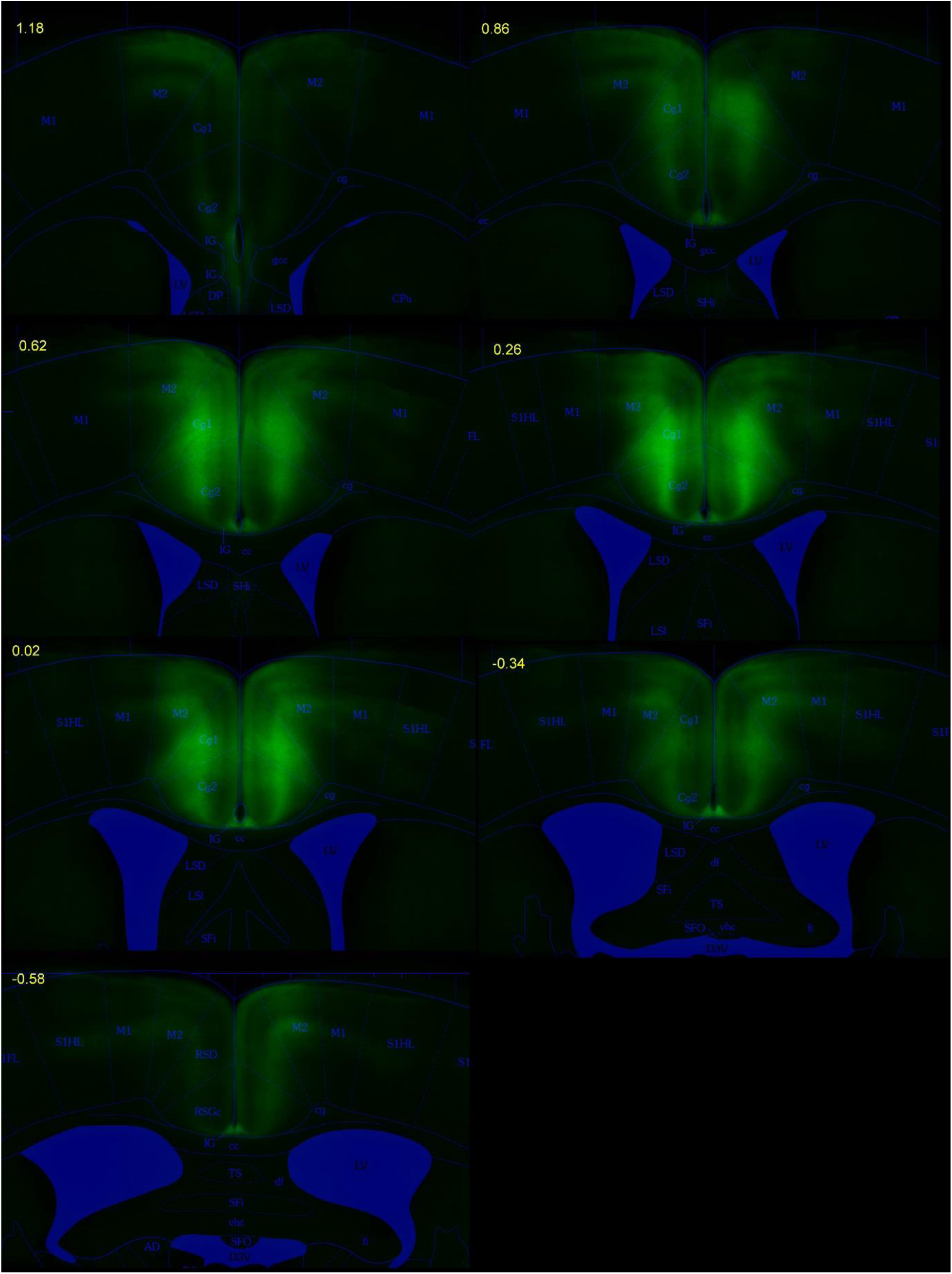
JAWS expression. Average JAWS-GFP fluorescence for all JAWS-GFP animals included in the study aligned onto reference atlas (Paxinos and Franklin, 2007). Numbers indicate anterior-posterior position relative to bregma (mm).

**Figure S5.**
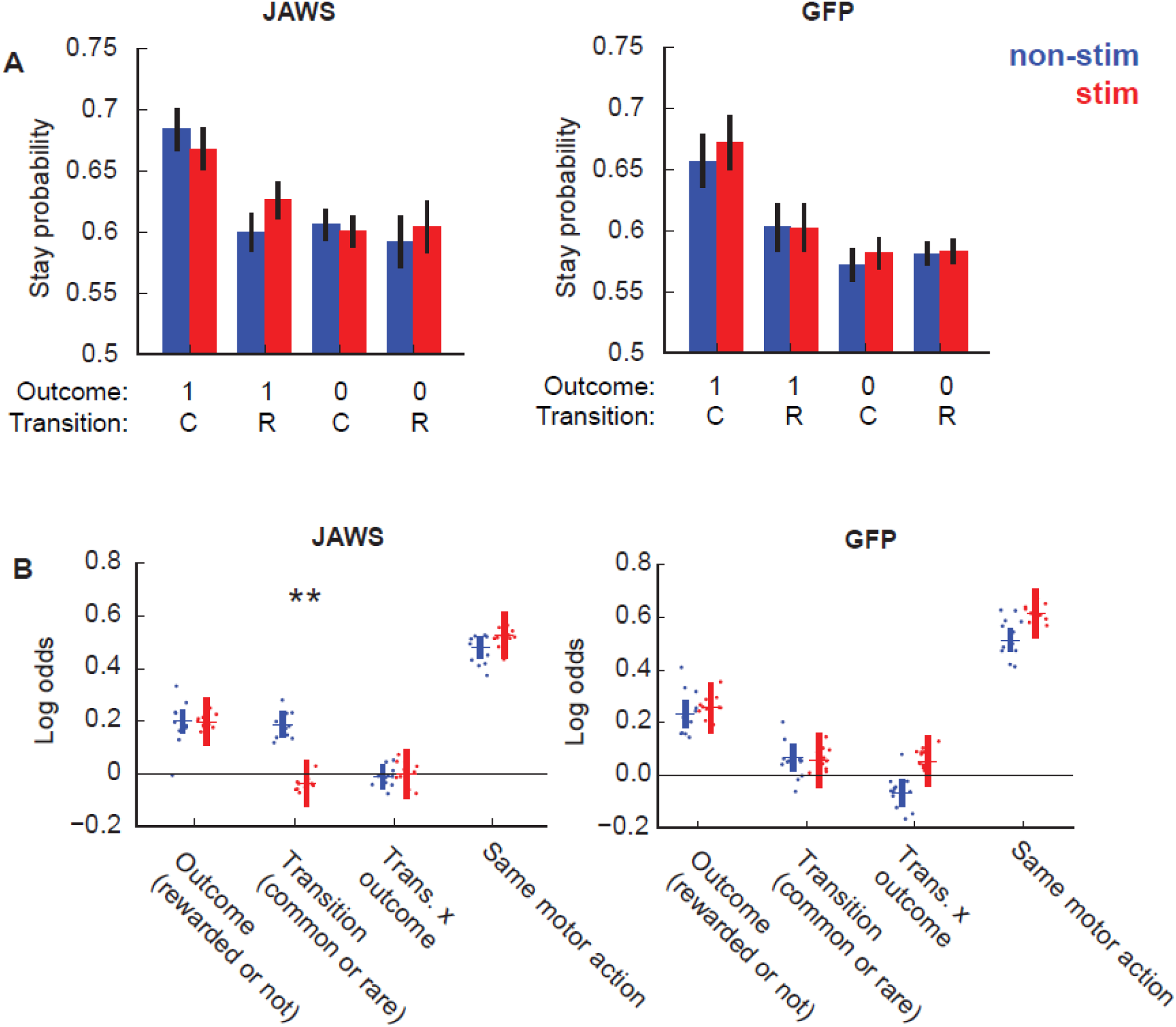
Optogenetic silencing of ACC in two-step task. **A)** Stay probabilities analysis on stimulated (red) and non-stimulated (blue) trials in JAWS (top panel) and GFP (bottom panel). **B)** Regression analysis including additional predictor *same motor action* – repeat choices if this requires the same motor action (e.g. left→top).

**Figure S6.**
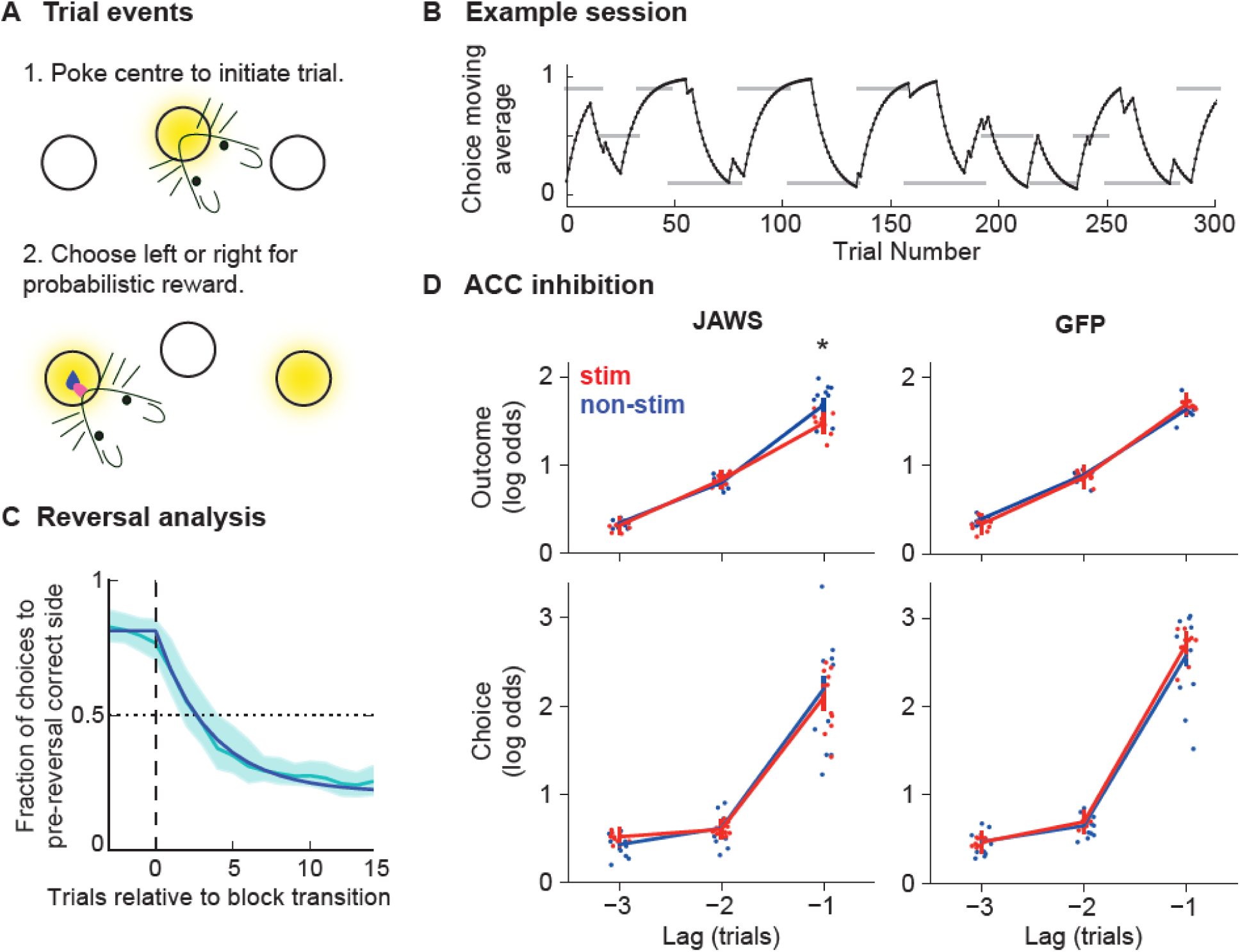
Optogenetic silencing of ACC in probabilistic reversal learning task. **A)** Diagram of apparatus and trial events. **B)** Example session, black line shows exponential moving average (tau = 8 trials) of choices, grey bars indicate reward probability blocks with y position of bar indicating whether left or right side has high reward probability or a neutral block. **C)** Choice probability trajectories around reversal in reward probabilities: Pale blue line – average trajectory, dark blue line – exponential fit, shaded area – cross-subject standard deviation. **D)** Logistic regression analysis showing predictor loadings for stimulated (red) and non-stimulated (blue) trials, for the ACC JAWS (left panel) and GFP controls (right panel). Bars indicate 95% confidence intervals on the population mean, dots indicate maximum a posteriori (MAP) subject fits. * indicates significant difference (Bonferroni corrected P < 0.05) between stimulated and non-stimulated trials.

**Figure S7.**
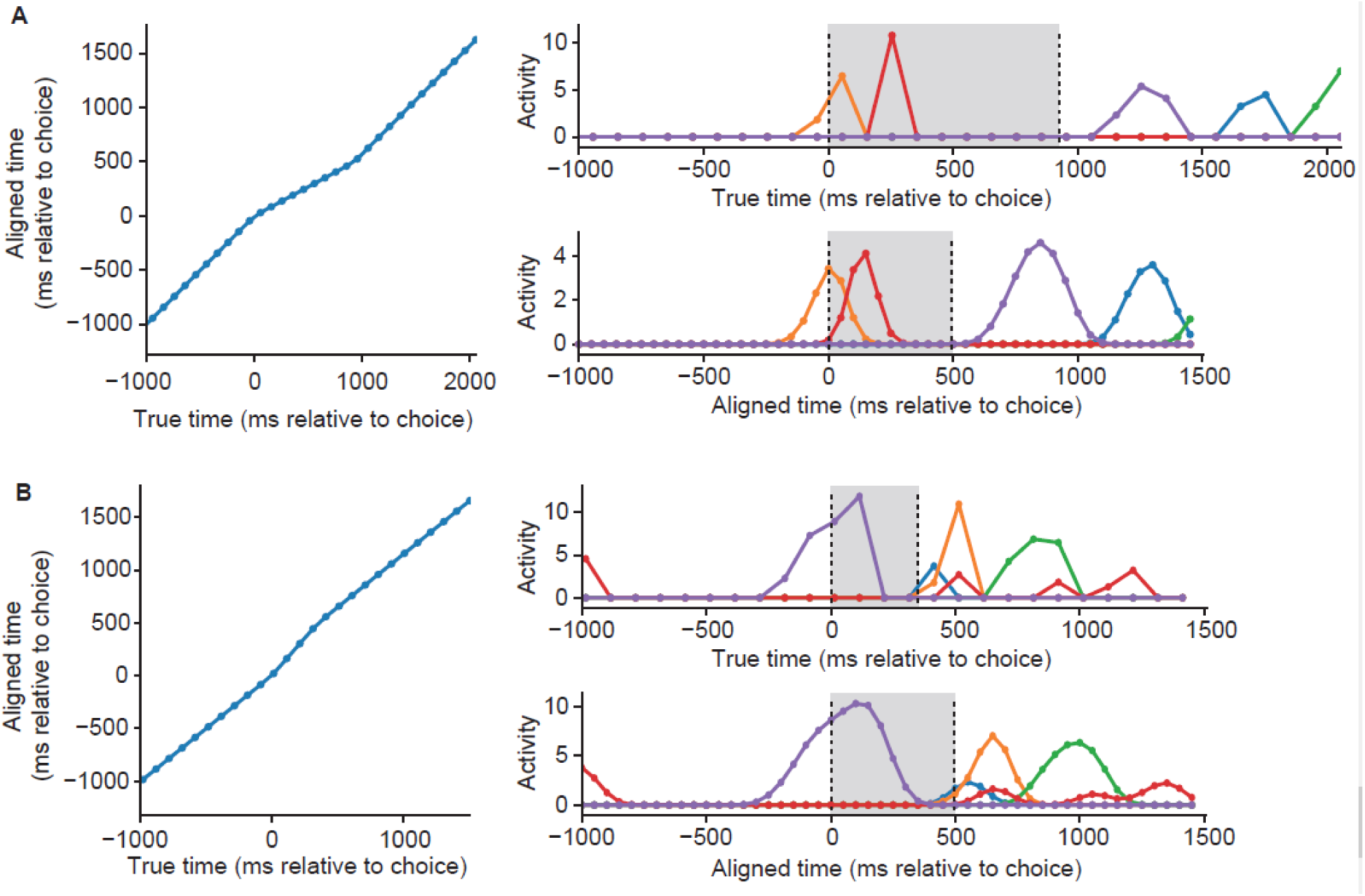
Calcium imaging alignment, up-sampling and smoothing. **A)** Alignment of imaging data on a trial where the interval between choice and second-step port entry was longer than the median interval. Left panel shows the true and aligned times of microscope frames plotted against each other. Right top panel shows the activity of 5 neurons before alignment. Vertical dashed lines show the times of choice and second-step port entry. Right bottom panel shows the activity of the same 5 neurons after alignment, up-sampling and smoothing. Grey shaded regions indicate the interval between choice and second-step port entry that is time-warped **B)** As for **A** but for a trial where the interval between choice and second-step port entry was shorter than the median interval.

## Supplementary Results

### Comparison of task variants with and without transition probability reversals

We introduced reversals in the transition probability mapping the first-step actions to the second-step states, because without them, extensively trained animals could in principle learn strategies that look like model-based RL but in fact rely on latent state inference rather than planning (Akam et al., 2015). We therefore asked what impact dynamically changing transition probabilities had on behaviour by running a version of the task where the transition probabilities linking the first step actions to second-step states were fixed (n=10 mice, 240 sessions analysed from day 22+ of training). Subjects were much better at tracking which option was best, choosing the correct option at the end of blocks on 0.83 ± 0.04 (mean + SD) of trials, and adapting to reversals with a time constant of 6.5 trials (P < 0.001 for difference between tasks on both measures, permutation test) (Figure S3A). Note that fixing the transition probabilities does not change the contrast between good and bad choices in terms of their reward probabilities. The granular structure of behaviour was also different (Figure S3 B-D), with a very strong influence of the transition-outcome interaction on the subsequent choice (P < 0.001, bootstrap test), a strong influence of the state transition (P < 0.001), but no direct influence of the trial outcome (P = 0.42) (between task differences at trial -1: P < 0.001 for stronger loading on transition and transition-outcome interaction predictor, P = 0.031 for weaker loading on outcome, permutation test).

These data show that in the fixed task, where subjects can, in principle, learn habit-like mappings from where rewards have recently been obtained to the correct first-step action (e.g. rewards on the left → choose up), overall performance was higher and behaviour showed a strong transition-outcome interaction, which can be generated by model-based RL or such latent state inference based strategies (Akam et al., 2015). The striking differences between behaviour on the fixed task and the version with transition reversals suggest that subjects do indeed solve them using different strategies. As our aim is to address neural mechanisms of model-based planning, for our investigation of ACC we focussed on the version of the task with changing transition probabilities designed to resist latent state strategies.

### Model comparison

The starting point for our model comparison was the RL agent used in the Daw two-step task (Daw et al., 2011). As the action-state transition probabilities in our task were not fixed, we modified the model-based component of the agent to update its estimate of the transition probabilities for the chosen action on each trial using an error driven learning rule. As in the original Daw agent we included a perseveration parameter which promoted repeating the previous choice.

We observed that some subjects appeared to have a bias to move either clockwise or counter-clockwise around the set of pokes (e.g. left→top, right→bottom). Including this predictor in the logistic regression model substantially improved the models integrated Bayes Information Criterion (Δ iBIC = 2639). Subjects may have developed these biases because it is the simplest fixed response pattern that was not penalised by the block transition rule (as block transitions were triggered based behaviour, a bias for the top or bottom port resulted in that port spending more the time as the bad option). Based on the evidence for this ‘rotational’ bias in the logistic regression, we included it in the RL models in addition to a standard choice bias.

We compared the goodness of fit of a pure model-free agent, a pure model-based agent, and an agent which used a mixture of both strategies. The mixture agent provided a better fit to the data than either the pure model-free (Δ iBIC = 264, Figure S2A) or pure model-based agent (Δ iBIC = 888), and the mixture model fit suggested an approximately equal contribution of model-based and model-free control. However, as the task is novel and hence we do not know what features may be present in the behaviour, we performed an exploratory process of model comparison to test whether adding additional features better captured the behaviour. This identified a number of features which greatly improved fit quality.

RL models typically assume that values of actions that are not chosen remain unchanged. However, it has been reported that model-fits in some rodent decision making tasks are substantially improved by including forgetting about the value of not chosen actions, typically implemented as action value decay towards zero (Ito and Doya, 2009, 2015). Including such action value forgetting in the mixture agent produced a dramatic improvement in iBIC score for our data (Δ iBIC = 7698). Including forgetting about action-state transition probabilities, implemented as a decay of transition probabilities for the not chosen action towards a uniform distribution, further improved the goodness of fit (Δ iBIC = 643). The mixture agent including value and transition probability forgetting again showed approximately equal weighting of the model-based and model-free action values in controlling behaviour. When forgetting was included for each agent the mixture agent provided a better fit to the data than either a pure model-free (Δ iBIC = 612) or pure model-based (Δ iBIC = 3066) agent.

Forgetting decreases the value of not chosen relative to chosen options, and therefore promotes choice perseveration. It is therefore possible that if subjects are in fact strongly perseverative, this could be mistakenly identified as forgetting. Though the model included a perseveration parameter for repeating the previous choice, several studies have reported perseveration effects spanning multiple trials, even in tasks where decisions optimally should be treated as independent (Gold et al., 2008; Akaishi et al., 2014). We therefore tested whether goodness of fit was improved by an exponential choice kernel through which prior choices directly influenced the current choice with exponentially decreasing weight at increasing lag. This is equivalent to the decision inertia model of Akaishi et al. (2014). The addition of this exponential choice kernel dramatically improved fit quality when added to the mixture agent without forgetting (Δ iBIC = 7133). However even with the exponential choice kernel included, value forgetting substantially improved goodness of fit (Δ iBIC = 2071), and transition probability forgetting further increased goodness of fit (Δ iBIC = 194). These results indicate that forgetting about values and transitions for not chosen options is a genuine feature of the behaviour and not an artefact due to perseveration. They further indicate that subjects do in fact show a strong tendency to perseverate over multiple trials, which is not captured even by forgetting RL models, presumably because it is independent of the recent reinforcement history. Forgetting may be a heuristic used in dynamic environments where evidence becomes less reliable with the passage of time due to state of the world changing. Alternatively, forgetting may occur due to limitations of the learning systems involved, perhaps due to discrepancy between the rapidly changing reward statistics in the task and those typical of natural environments.

The choice kernel assumes that perseveration occurs at the level of the decision between the top and bottom pokes. However, in the current task, a given choice (top or bottom) entails a different motor action depending on which side (left or right) the previous trial ended on. We therefore considered a model with perseveration at the motor level such that the choice on a given trial only increased the probability of repeating the same motor action in future, e.g. a choice taken by moving from the left to top poke only increased the probability of choosing top in future following trials which ended on the left side. Motor perseveration was modelled by maintaining separate moving averages of choices following trials that ended on the left and right, which each influenced choices following trials ending on their respective sides. Replacing the exponential choice kernel with this motor level perseveration substantially improved fit quality (Δ iBIC = 1004). However, including perseveration both at the level of choice, (top vs bottom, independent of motor action), and at the motor level, further improved fit quality (Δ iBIC = 499), indicating that subjects have perseverative tendencies at both choice and motor levels that are not predicted by the RL component of the model. These data support the existence of mechanisms which reinforce selected behaviours in a reward-independent fashion, i.e. simply choosing to execute a behaviour increases the chance that behaviour will be executed in future. This is consistent with previous reports from perceptual (Gold et al., 2008; Akaishi et al., 2014) and reward-guided decision making tasks (Miller et al., 2019), and we think is a parsimonious explanation for our results. Such perseveration may be a signature of a mechanism for automatizing behaviour by reinforcing chosen actions. Thorndike proposed such a ‘law of exercise’ (1911) and the idea has recently been revisited by Miller et al. (Miller et al., 2019) who suggest that habit formation occurs through outcome-independent reinforcement of chosen actions. This framework views habit formation as a supervised learning process in which behaviour generated by value sensitive systems, i.e. model-free and model-based RL, is used to train value-independent learning systems. Such a mechanism could account for multi-trial perseveration effects observed in our data. An alternative mechanism which could generate perseveration would be subjects sampling an option multiple times between choices, which may be adaptive if the decision process is costly in time or effort. However, this does not account for the observation in our data that perseveration occurred at the level both of choices and of motor actions, with different timescales for each (see respective learning rates, Figure S2 C).

Evidence that perseveration occurred both at the level of choice and motor action raises the question of whether reward driven learning also occurs at both levels of representation. This might be expected from the architecture of parallel cortical-basal ganglia loops, with circuits linking somatosensory and motor cortices to dorsolateral striatum learning values over low level motor representations, and circuits linking higher level cortical regions to medial and ventral striatum learning values over more abstract state and action representations. Indeed, human two-step task behaviour shows evidence of model-free value accruing to low level sensory-motor features (Shahar et al., 2019). We therefore tested an agent in which model-free action values were learned in parallel for actions represented both in terms of choice (top/bottom) and motor action (e.g. left→top). This improved goodness of fit (Δ iBIC = 117) and the resulting model fit indicated that motor-level model-free values had a somewhat stronger influence on behaviour than the choice level model-free values. With multi-trial perseveration kernels and motor level effects included in each model, the mixture agent again provided a better fit to the data than either a pure model-free (Δ iBIC = 127) or pure model-based (Δ iBIC = 227) agent.

We tested a number of other modifications to the model including separate learning rates at the first and second step, but did not find further improvements in fit quality (Figure S2B). Finally, as adding features to the model may make other features which previously improved the fit unnecessary, we tested whether removing any individual component from the model improved fit quality but again did not find further improvements (Figure S2B).

### Motor effects do not explain ACC inhibition effect on transition predictor

Evidence for perseveration and model-free RL at the motor level raises a possible alternative interpretation of why ACC inhibition reduced the influence of common vs rare state transitions on choices. This is because the state transition determines which second-step state the subject ends up in, and hence the motor action required to repeat the choice on the next trial. To test whether motor-level factors can account for the ACC inhibition effect, we analysed the ACC inhibition data using a logistic regression analysis including an additional predictor which coded a tendency to repeat choices when this required the same motor action as the previous trial (Figure S5B). Although *same motor action* significantly predicted repeating choice (P < 0.0001, bootstrap test), ACC inhibition had no effect on the *same motor action* predictor (P = 0.94 uncorrected), and the effect of ACC inhibition on the common/rare transition predictor remained significant (Bonferoni corrected P = 0.0032, stim-by-group interaction P = 0.032). We also tested whether the observed correlation between the ACC inhibition effect on the transition predictor and subjects use of model-based RL (Figure 6E) was specific, by using a multiple linear regression to predict the strength of opto effect across subjects using a set of parameters from the RL model: model-based weight (*G*_*mb*_), model-free weight (*G*_*mf*_), motor model-free weight (*G*_*mo*_), and motor-perseveration (*P*_*m*_). Model-based weight predicted the strength of opto effect on the transition predictor (P = 0.03), but none of the other parameters did (P > 0.45). Together these results argue that the effect of ACC inhibition on sensitivity to action-state transitions is mediated by disrupted model-based RL and not motor level factors.

### ACC inhibition in a probabilistic reversal leaning task

We assessed the effects of the same ACC manipulation used in the two-step task on a probabilistic reversal learning task (n = 10 JAWS mice, 202 sessions, 10 GFP mice, 202 sessions). In this task both model-free and model-based RL are expected to generate qualitatively similar influence of trial events on subsequent choice, i.e. rewarded choices will be reinforced, though there may be quantitative differences if the model-based system is able to learn the block structure and infer block transitions rather than relying on TD value updates.

Subjects initiated trials in a central port, then chose left or right for a probabilistic reward (Figure S6A). Mice tracked the correct option (Figure S6 B,C), choosing correctly at the ends of blocks with probability 0.80 ± 0.04 (mean ± SD), and adapting to reversals with a time constant of 3.57 trials (exponential fit tau). Parameters for optogenetic silencing were matched as closely as possible to those used in the two-step task, with the same viral vector, injection sites and stimulation parameters. Stimulation was delivered from when subjects poked in the side port and received the trial outcome until the subsequent choice.

We assessed the effect of ACC silencing using a logistic regression analysis with previous choices and outcomes as predictors (Figure S6 D). Previous choices predicted the current choice with decreasing influence at increasing lag. Rewards predicted repeating the rewarded choice, with decreasing influence at increasing lag. ACC inhibition subtly reduced the influence of the most recent outcome (permutation test P = 0.024 Bonferroni corrected for 6 predictors, stimulation-by-group interaction P = 0.014). These data suggest that while ACC did participate in this simple reward guided decision task, its contribution could largely be compensated for by other regions, consistent with model-based and model-free control both recommending repeating rewarded choices.

## Notes

### Competing Interest Statement

The authors have declared no competing interest.

### Summary of Updates

- Added two-step task calcium imaging data. - Added additional analysis showing correlation between opto effect on transition predictor and subjects use of model-based RL.

